# Unfolding and Translocation of Knotted Proteins by Clp Biological Nanomachines: Synergistic Contribution of Primary Sequence and Topology Revealed by Molecular Dynamics Simulations

**DOI:** 10.1101/2021.04.30.442167

**Authors:** Hewafonsekage Yasan Y. Fonseka, Alex Javidi, Luiz F. L. Oliveira, Cristian Micheletti, George Stan

## Abstract

We use Langevin dynamics simulations to model, at atomistic resolution, how various natively–knotted proteins are unfolded in repeated allosteric translocating cycles of the ClpY ATPase. We consider proteins representative of different topologies, from the simplest knot (trefoil 3_1_), to the three–twist 5_2_ knot, to the most complex stevedore, 6_1_, knot. We harness the atomistic detail of the simulations to address aspects that have so far remained largely unexplored, such as sequence–dependent effects on the ruggedness of the landscape traversed during knot sliding. Our simulations reveal the combined effect on translocation of the knotted protein structure, i.e. backbone topology and geometry, and primary sequence, i.e. side chain size and interactions, and show that the latter can even dominate translocation hindrance. In addition, we observe that, due to the interplay between the knotted topology and intramolecular contacts, the transmission of tension along the peptide chain occurs very differently from homopolymers. Finally, by considering native and non–native interactions, we examine how the disruption or formation of such contacts can affect the translocation processivity and concomitantly create multiple unfolding pathways with very different activation barriers.

## Introduction

Controlled protein degradation to remove excess or non–functional proteins is an essential regulatory mechanism that ensures cell survival upon various stresses, maintains protein homeostasis and prevents pathogenic pathways. ^1–3^ Various enzymes of the AAA+ superfamily, such as bacterial caseinolytic protease (Clp) ATPases and the eukaryotic proteasome, assist the mechanical degradation process of substrate proteins (SPs) by actively threading them through a narrow pore,^4^ causing their mechanical unfolding. Most frequently, translocation initiates at one of the protein termini, which is engaged through a covalently attached tag or an intrinsic recognition sequence. ^5–7^ These non–specific recognition mechanisms are key for protein quality control as they allow Clp ATPases to act promiscuously by processing diverse substrates tagged for degradation. *In vitro* experiments^8–12^ have given support to the notion that knotted proteins too can undergo *in vivo* proteolysis via Clp–mediated degradation.

The repeated pulling action exerted locally by central channel loops of ATPases can efficiently unravel protein domains with large global stability, such as malate dehydrogenase, dihydrofolate reductase or titin I27. ^13^ The resulting unfolding pathways can strongly depend on intrinsic properties of SPs, such as their secondary and tertiary content, and on the detailed interactions with ATPases. For instance, *β*–sheets generally present larger unfolding energy barriers than *α*–helical elements, but these differences can be significantly modulated by the pulling direction, e.g. *β*–sheets in the direction perpendicular or parallel to the hydrogen bond registry.^14–20^

Plasticity of the ATPase surface during allosteric cycle facilitates the re–orientation of the translocating SPs and can thus guide dynamic selection of the force application towards directions that are more favorable and conducive to unfolding.^21^ This mechanism is so critical for ATPase–mediated degradation that significant increases in unfolding resistance are observed when SPs’ freedom to re–orient is hindered by an externally applied force at the free terminal or by the presence of multiple folded domains, as in the case of tandem constructs. ^22^

Degradation of proteins with topological entanglement, such as knots, is of particular interest as it inherently requires overcoming distinctive free–energy barriers and novel unfolding and translocation mechanisms. The presence of molecular knots in proteins is evolutionarily intriguing given their potential to induce misfolding and disrupt folding pathways. ^23–25^ Nevertheless, physical knots have been identified in approximately 1% of protein data bank (PDB) entries and other forms of structural entanglements were found in approximately 6% of them.^26–32^ So far, four distinct knot types have been algorithmically detected in proteins, namely trefoil, 3_1_, figure–of–eight, 4_1_, three–twist, 5_2_, and stevedore, 6_1_, knots.^28^ Multiple computational studies highlighted details of folding pathways leading to knot formation, including the role played by non–native interactions. ^33–41^ Structural^28^ and phylogenetic evidence^25^ suggests that several instances of knotted proteins have evolved from unknotted ones, though the specific functional advantage afforded by knots is still unclear.^42^ It has long been debated whether knots can enhance thermodynamic stability^43^ or the mechanical one. Force spectroscopy experiments have revealed a strong dependence of the stretching response on the pulling direction. When the stretching force is applied along the N–C direction, the mechanical response is modulated by the progressive tightening of the knot. ^44–49^ However, pulling in other directions can elicit very different responses associated to transitions to a different knotted state, or even the complete unknotting, as it was established for the 5_2_–knotted protein UCH–L1.^48^

Experimental and computational studies have further shown that knots may affect the translocation of knotted proteins and general polymers through pores that are too narrow to allow for the simultaneous passage of multiple strands. ^28,50–56^ The progressive tightening of the knot at the pore entrance brings the amino acids (or monomers) of the knotted region into close contact and such interactions create a rugged energy landscape that the chain has to negotiate to slide along its contour. The height of the barriers, which depends on knot topology, ^54^ increases with the driving force and when the latter is sufficiently large, the translocation process can come to a stall. ^51^

The depth of knot can affect the translocation process too. Proteins with shallow knots, in which the knot boundary lies within 30 residues of the free protein terminal, have a greater probability of successful translocation as the knot can spontaneously slip off the chain. ^8,56^ Recent experimental studies have shown that ClpXP can completely degrade the 3_1_–knotted protein MJ0366 of *Methanocaldococcus jannaschii*, which contains a shallow knot. ^8^ Interestingly, ClpXP is also able to completely degrade the 3_1_–knotted YbeA methyltransferase of *Escherichia coli*, which contains a deep knot.^9^ In both cases, however, the degradation process is stalled by addition at the distal terminal of stable domains, such as the green fluorescent protein or ThiS, to yield partial degradation fragments. Additional modulation of degradation can be provided by the sequence directionality of substrate processing. Intriguingly, for the 5_2_–knotted protein UCH–L1, efficient degradation was observed for processing in the N–C direction,^9^ whereas strong degradation resistance is manifest in the C–N direction.^9–12^ The enhanced resistance for C–terminal pore entries has so far been rationalized in terms of the different local mechanical properties^9^ and knot depth with respect to the N terminus.^11,12^

Coarse–grained models of unfolding and translocation of knotted proteins and polymers through solid–state nanopores provided important clues for degradation mechanisms. ^51–57^ For example, the translocation rate of a 5_2_–knotted polypeptide was found to be approximately 100 times lower than that of the unknotted counterpart. ^57^ Mechanical pulling of the 3_1_–knotted proteins YibK, YbeA, and MJ0366 and the 5_2_–knotted protein UCH–L3 through a narrow pore showed knot sliding and removal of the trefoil knots, but knot tightening and stalled translocation of the 5_2_ knot.^56^ Interestingly, twist knots, such as all those found in proteins, offer much more resistance to translocation than other knot types. ^54^

To date, however, the effect of repetitive conformational transitions of ATPase subunits and the role of non–native SP interactions on unfolding and translocation mechanisms of knotted proteins mediated by AAA+ nanomachines is not well understood.

In this study, we use molecular dynamics simulations to probe translocation and unfolding mechanisms of knotted SPs mediated by Clp ATPase biological nanomachines. To this end, we use atomistic Langevin dynamics to probe the unfolding and translocation of three knotted SPs with 3_1_, 5_2_ and 6_1_ knot types mediated by the ClpY nanomachine. Our analysis of the time–dependent evolution of native and non–native contacts, of knot sliding, SP’s re–orientation, translocation hindrance and tension propagation reveal a complex interplay of SPs’ structural architecture and primary sequence in determining the resulting Clp threading compliance.

## Methods

### Implicit Solvent Model of ClpY∆I and Knotted Substrate Proteins

We describe the ClpY∆I–SP system by using the EEF1 implicit solvent model,^58,59^ which enables us to study, at an atomistic level, the interactions between the ClpY ATPase and knotted SPs in a computationally efficient manner.

In this study, we use three knotted proteins, with diverse knot topology, namely 3_1_, 5_1_ and 6_1_ knots (Table 1). The protein with the simplest topology, a trefoil (3_1_) knot, is the MJ0366 protein of *Methanocaldococcus jannaschii*. Our simulations model the 82–residue segment, between sequence positions 6–87, resolved in the crystal structure corresponding to the Protein Data Bank (PDB) ID 2EFV^60^ (Figure 1A). The backbone of the same 2EFV entry was also used to design a MJ0366 variant without side chains. The resulting structure is thus a trefoil–knotted polyG 82–mer. For the 5_2_ knot topology we considered the solution structure of the S18Y mutant of the 231–residue ubiquitin carboxy–terminal hydrolase L1 (UCH–L1) from *Homo sapiens*, with PDB ID 2LEN^61^ (Figure 1B). For the third and most complex protein topology, corresponding to the 6_1_ knot, we considered the *α*–haloacid dehalogenase I from *Pseudomonas putida*. We modeled the 295–residue chain B, between sequence positions 2–296, of the crystal structure with PDB ID 3BJX^62^ (Figure 1C). An (SsrA)_2_ degradation tag (SsrA sequence AANDENYALAA) was covalently attached at the C–terminal and/or at the N–terminal of each knotted proteins, and provided the leading peptide to initiate translocation on the resulting fusion protein (Table 1). In our analysis, all amino acids of knotted proteins are numbered sequentially starting from 1 at the N terminal. During translocation simulations, the center of mass of the ClpY ATPase is maintained near the origin of a Cartesian reference system and the pore axis is aligned with the *z*–axis, which is oriented such that the *cis*, or proximal, side corresponds to negative values of *z* and the *trans*, or distal, side to positive ones. The SP is initially oriented with its principal axis of inertia aligned with the *z*–axis and its center of mass is located at *z* ≃ − 60 Å on the cis side of ClpY∆I ATPase. The (SsrA)_2_ peptide, which has an extended configuration, is partially inserted into the ClpY pore (Figure 1). To avoid systematic biases of the initial configurations, SPs are rotated through an arbitrary angle about the *z*–axis at the beginning of simulations. ClpY∆I mediated translocation and unfolding of the SPs are investigated using Langevin dynamics simulations at T = 300 K, with a friction coefficient of 5 ps^−1^ and a time step of 2 fs. Simulations are performed with the CHARMM simulation package^63^ using the Extreme Science and Engineering Discovery Environment (XSEDE) supercomputer resources. ^64^

**Table 1:** Summary of folded domains used in substrate proteins studied and number of trajectories performed in this work.

| Substrate protein<sup>a</sup> | Knot Type | Sequence length<sup>b</sup> | $\langle F \rangle$(pN) | N<sub>traj</sub><sup>c</sup> |
| --- | --- | --- | --- | --- |
| 2EFV-(SsrA)<sub>2</sub> | 3<sub>1</sub> | 82 | 300 | 28 |
| 2EFV-(SsrA)<sub>2</sub> | 3<sub>1</sub> | 82 | 200 | 20 |
| PolyG-(SsrA)<sub>2</sub> | 3<sub>1</sub> | 82 | 100 | 24 |
| 2LEN-(SsrA)<sub>2</sub> | 5<sub>2</sub> | 231 | 500 | 10 |
| (SsrA)<sub>2</sub>-2LEN-(SsrA)<sub>2</sub> | 5<sub>2</sub> | 231 | 500 | 7 |
| (SsrA)<sub>2</sub>-2LEN | 5<sub>2</sub> | 231 | 500 | 8 |
| 3BJX-(SsrA)<sub>2</sub> | 6<sub>1</sub> | 295 | 500 | 12 |
| I27-(SsrA)<sub>2</sub> | — | 98 | 200 | 20 |

<sup>a</sup> Knotted domains are identified according to the corresponding PDB ID. Substrate proteins are processed by the Clp ATPase in the C–N direction, except for the N-terminal tagged (SsrA)<sub>2</sub>-2LEN.
<sup>b</sup> Polypeptide sequence length of folded domains modeled in our simulations.
<sup>c</sup> The number of simulation trajectories performed at $T = 300$ K.

**Figure 1:**
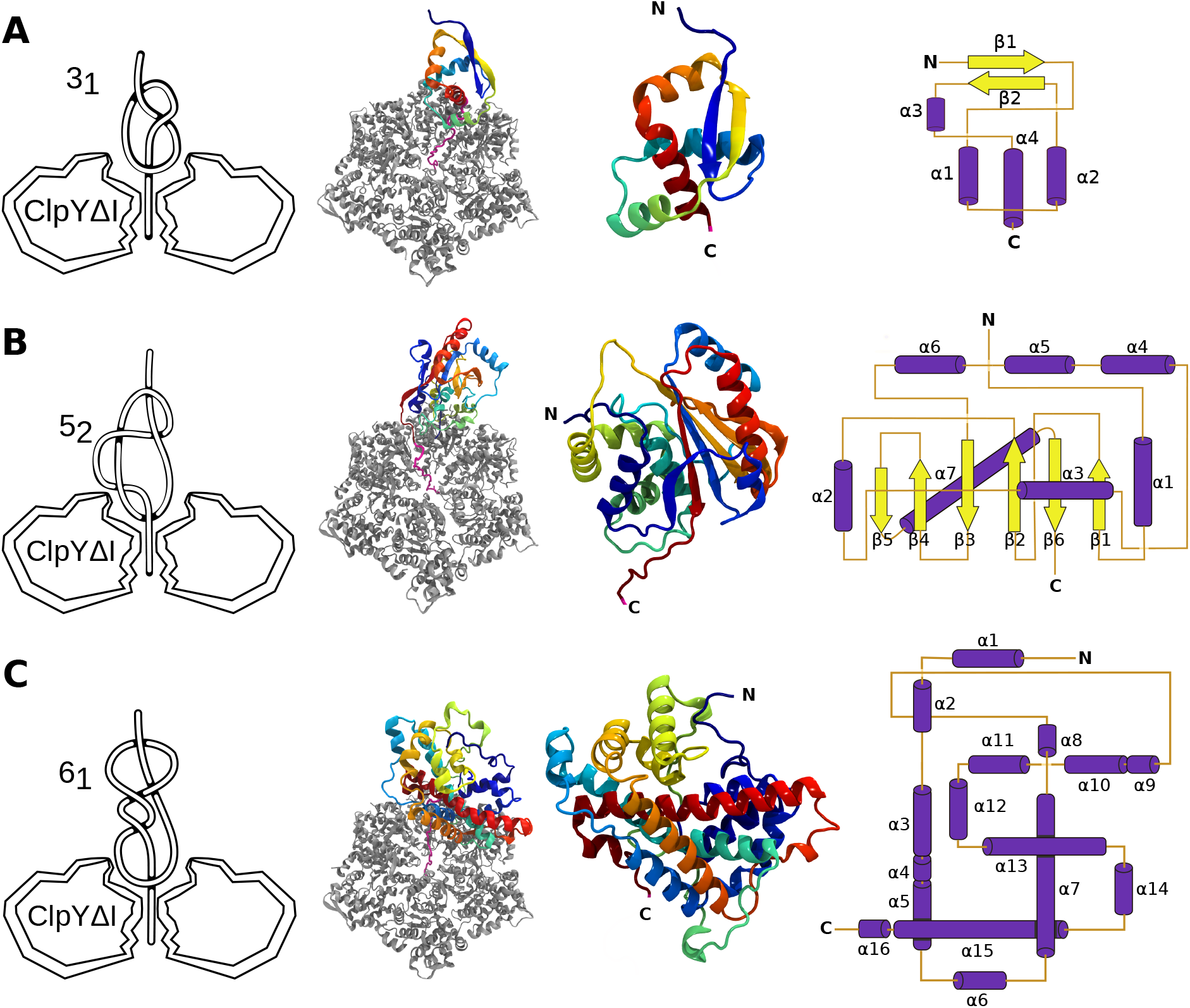
Remodeling of knotted proteins by ClpY∆I. Schematic view (left panels) and molecular representation (middle panels) of the simulation setup of the ClpY∆I (grey)–knotted proteins (color gradient from N–terminal, blue, to C–terminal, red), and native structure and connectivity maps (right panels) of (A) 3_1_–knotted MJ0366; (B) 5_2_–knotted ubiquitin carboxy–terminal hydrolase L1 (UCH–L1) and (C) 6_1_–knotted *α*–haloacid dehalogenase I substrate proteins. The (SsrA)_2_ recognition tag is covalently attached at the N– and/or C–terminals. Molecular structures are rendered using Visual Molecular Dynamics.^65^

### ClpY∆I Allosteric Motions

Our computational model incorporates fundamental aspects of the allosteric cycle of the Clp ATPase, namely repetitive binding and release of the SP in high and low affinity states. These are represented here by using the available crystal structures of *Escherichia coli* ClpY, namely PDB IDs 1DO2 (open pore) and 1DO0 (closed pore).^66^ The auxiliary I–domains of ClpY subunits (residues 111–242) are not included in our simulations, allowing us to probe common mechanisms of Clp ATPases. Transitions in the ClpY allosteric cycle are described by using the targeted molecular dynamics approach, which minimizes the root–mean–square deviation from the target (open or closed) state of the pore.^67^ Each hemicycle consists of six sequential transitions of individual ClpY subunits, represented by two chains, which undergo conformational changes between their open and closed states. The subunit undergoing the first transition in the hemicycle is randomly selected and subsequent transitions follow the clockwise ring order as viewed from the cis side of ClpY. In each step, the centers of mass of each of the five subunits not involved in the transitions are constrained to their current position. The total duration of each cycle is *τ* = 120 ps, which corresponds to an effective pulling speed of 1 Å/ps considering the 10 Å excursion of pore loops. As shown in our prior studies of ClpY–mediated threading of I27 SPs, simulations using this effective pulling speed yield results consistent with those obtained in simulations performed with lower speeds (0.2 Å/ps and 0.02 Å/ps)^22^ as well as with atomistic simulations of bulk mechanical unfolding.^68^ We also note that time scales probed in simulations using continuum models, such as implicit solvent, correspond to longer effective biological times. ^69–71^

### External Repetitive Force Coupled with Allosteric Motions

We use an external pulling force to accelerate SPs’ unfolding and translocation. The force is applied during the open → closed hemicycle to the SP segment that occupies the ClpY pore at the beginning of each single subunit transition. The force is directed along the pore axis (*z*–axis) and is distributed uniformly among SP’s backbone heavy atoms whose *z* coordinate satisfies |*z* − 〈*z*_loop_〉| < 5 Å, where 〈*z*_loop_〉 represents the average *z* coordinate of Tyr91 amino acids of central channel loops of ClpY at the beginning of each cycle. The magnitude of the driving force is drawn from a Gaussian distribution centered at 100 pN, for the 3_1_–knotted polyG SP, 300 pN, for the 3_1_–knotted MJ0366 protein, and 500 pN, for the 5_2_– and the 6_1_–knotted SPs. The standard deviation is set to 15 pN for the 3_1_–knotted cases and 30 pN for the 5_2_– and 6_1_–knotted SPs (Table 1). The number of backbone heavy atoms occupying the ClpY pore is about 20 atoms, therefore the applied force per atom has an average of ≃ 15 pN for 3_1_ knot and 25 pN for 5_2_ and 6_1_ knot types. We use lower forces for the 3_1_ polyG case due to its weak structural connectivity and larger forces for the 5_2_ and 6_1_ knots due to their strong structural connectivity. Compared to MJ0366, the more complex and larger proteins have a much slower translocation. We thus accelerated it at times larger than 1000*τ* by increasing the average force to 600pN.

We note that our pulling protocol uses forces that are several–fold larger than typically accessible to molecular motors, which are in the 50–100pN range.^72^ This is necessary to bridge the characteristic translocation times, with those accessible computationally. The use of large forces is customary in MD simulations of externally–driven biomolecules. It is justified by the typically sharp free–energy barriers that hinder the response of these systems to external driving, which result in a weak dependence of the transition state placement on the magnitude of the applied force. We have explicitly verified that this holds for our system by comparing the unfolding by translocation pathways of MJ0366 at the reference average force of 300 pN and at the lower one of 200 pN, see Supplementary Text and Fig. S1.

### Fraction of Native and Non–Native Contacts

The fraction of native contacts (Q_N_) is computed as 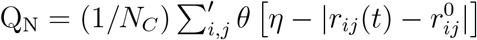, where N_C_ is the number of native contacts, *r*_*ij*_(*t*) is the minimum distance, at time *t*, between any two heavy atoms of residues *i* and *j* and 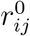 is the corresponding native distance. The primed summation symbol indicates that the sum is taken over residues at a chemical distance larger than 2, |*i* − *j*| > 2. The interaction cutoff distance for amino acids in the native state is set to 6 Å. *θ*(*x*) is the Heaviside step function for which *θ*(x) = 1 if x ⩾ 0 and *θ*(x) = 0 if x < 0, with tolerance *η* = 2 Å. The fraction of non–native contacts (f_NN_) is then defined as f_NN_ = N_NC_/N_C_, where N_NC_ is the number of non–native residue pairs identified using the 6 Å cutoff.

### Evolution of the Translocated Fraction and Waiting Times

The progress of translocation is quantified through the time evolution of the translocated fraction, *x*(*t*), which is the instantaneous fraction of amino acids that have moved to the *trans* side, such that the axial location of the *C*_*α*_ atom satisfies *z*_*i*_(*t*) > *z*_*trans*_ = 12 Å. This threshold value of the *z* coordinate was chosen because it lies beyond the maximum *z* excursion of the ClpY pore loops. At each time *t*, the sequence position I of the most recently translocated amino acid is identified by using *z*_*I*_(*t*) = *min*{*z*_*i,trans*_(*t*)}.

In addition, we monitor the so–called waiting time,^73^ *w* = *dt/dx*, which is evaluated as follows. The translocated chain fraction, *x*, is subdivided in equal intervals of width ∆, and the ensemble of *x*(*t*) curves is used to compute the average translocation time for each interval. The set of discrete average times is then interpolated using the Savitzky–Golay (SG) algorithm^74^ and the waiting times are finally computed using finite differences for the interpolated translocation times. For the 3_1_ knotted SP, we use ∆ = 0.05 and the SG interpolation with the window length Γ_*SG*_ = 11 and the polynomial of order *m* = 2. For the polyG SP, ∆ = 0.035, Γ_*SG*_ = 13 and *m* = 3, and for I27, ∆ = 0.04, Γ_*SG*_ = 15 and *m* = 3. The waiting time of residue I is estimated as its residence time in the vicinity of the translocation boundary such that *z*_*I*_(*t*) ⪆ *z*_*trans*_.

### Detection of the knotted region

The evolution of the protein region hosting the knot was characterized with the Kymoknot webserver.^75^ Knot boundaries are denoted as k_N_ and k_C_ according to their greater proximity to the N and C terminals, respectively.

## Results

### Knot Tightening and Sliding Results in the Formation of Non–Native Contacts in the 3_1_–Knotted Substrate Protein

First, we consider the translocation of the MJ0366 SP, which features a shallow trefoil knot, bounded by 10 N–terminal residues and 9 C–terminal residues, that ensnares all of the secondary structural elements.^25,28,35,76,77^ In our simulation model of the nonconcerted ATP–driven allosteric cycles of ClpY∆I (see Methods, Figure 1A and Table 1), the SP is pulled through the ClpY central channel by stochastic axial forces exerted by channel loops and by an additional external force. The latter is introduced to accelerate the unfolding and translocation processes and make them amenable to atomistic simulations (see Methods).

As the SP C–terminal is engaged first, these simulations feature mechanical unfolding events, polypeptide translocation proceeding with C–N directionality. In all 28 recorded trajectories the knot tightens and slides to the N–terminal over time scales of order 1000 *τ*, hence comparable to the duration of the simulations (Supporting Information Movies S1 and S2).

Detailed analysis reveals a two–fold kinetic partitioning of the process. Over the simulated time scales, 50% (14) of translocations reach completion, with the knot sliding off the chain. In the remaining 50% (14) of cases, the chain remains knotted and only a partial SP translocation occurs (Figure 2). As shown in Figures 2A–B, in the first pathway, with a lower energy barrier, we observe a rapid decrease of the fraction of native contacts, Q_N_, with most native contacts lost on a time scale of ≃ 100 *τ*, and concomitant formation of non–native contacts, with f_NN_ ≃ 0.35 on this time scale. In the second pathway, with a higher energy barrier, we note the larger dispersion and slower decay of the fraction of native contacts, complemented by a significant fraction of non–native contacts, f_NN_ ≃ 0.25 over the entire simulation duration, ∼ 1000 *τ* (see Methods and Figures 2E–F). Strikingly, divergent types of interactions contribute to native and non–native contacts present during the Clp–mediated remodeling of the SP. As shown in Figures 2A,E, after ≃ 100*τ*, the decaying fraction of native contacts comprises balanced contributions of backbone–backbone interactions, which stabilize secondary structure elements, and side chain–side chain or side chain–backbone interactions, which support tertiary contacts. By contrast, the larger fraction of non–native contacts is dominated by side chain–based interactions (Figures 2B,F). These results indicate that the process of knot sliding towards the free terminal, rather than being well–described by smooth gliding over the unentangled backbone segment, is more aptly illustrated by traversing a rough conformational landscape. In the low energy barrier pathway, the mean unknotting time is 〈*t*_1_〉 = 451 ± 36 *τ*, and the SP completely translocates with knot tightening to a minimum length of 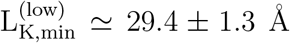 (Figures 2C–D). By contrast, as shown in Figures 2G–H, in the high energy barrier pathway the knot tightens more strongly to 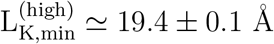, and translocation does not reach completion.

**Figure 2:**
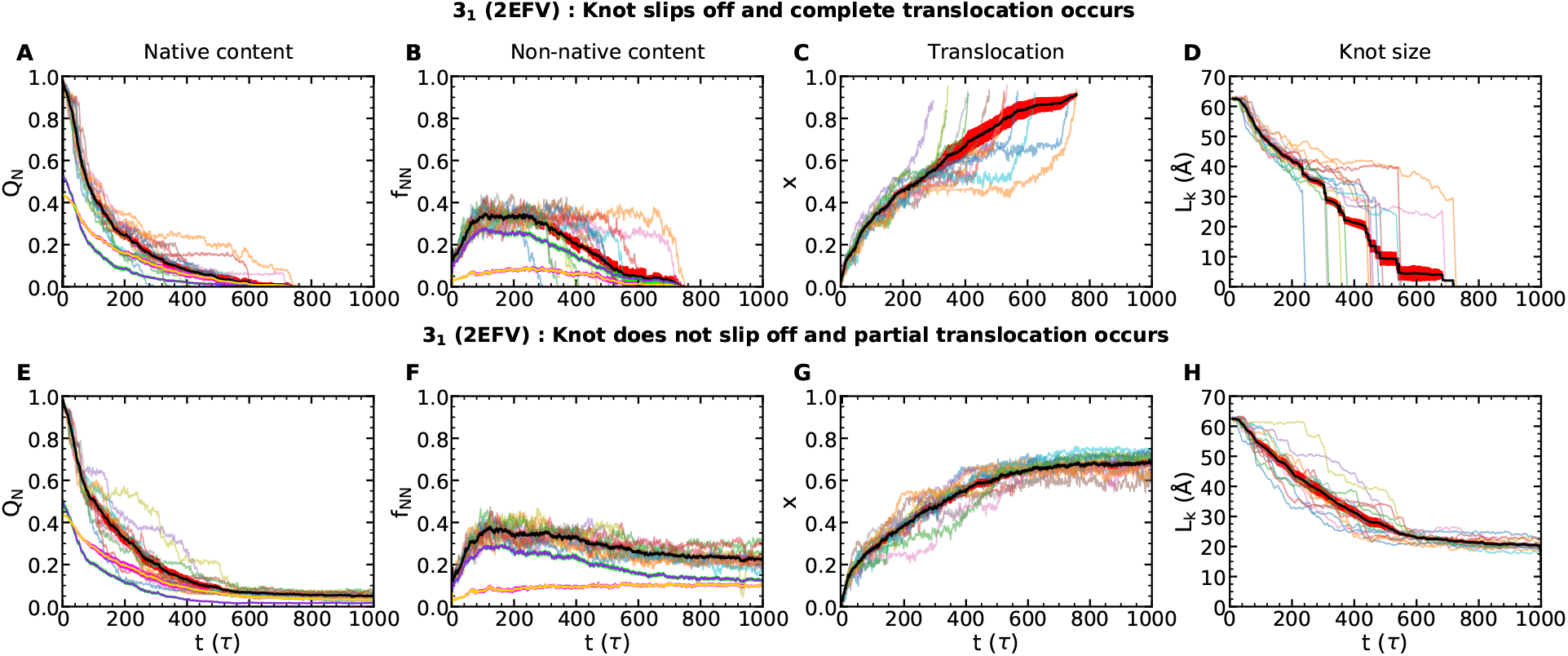
Kinetic partitioning of the 3_1_–knotted SP. Time evolution of the fraction of native contacts Q_N_, the fraction of non–native contacts f_NN_, the translocated fraction *x* and the knot length L_K_ in trajectories in which (A–D) the knot slips off the chain and the SP is completely translocated or (E–H) the knot does not slip off and translocation becomes stalled. Individual traces (thin curves) and averages and standard errors (thick curves; black and red, respectively) are shown. Average contribution (and standard errors) of contacts involving backbone–backbone interactions (yellow and red) and side chain–side chain or side chain–backbone interactions (purple and cyan) are also indicated.

To glean the molecular details that characterize these pathways, we analyzed the time evolution of the fraction of native and non–native contacts of each secondary structure element (Figures 3A–B). In the low energy barrier pathway, native contacts of secondary structural elements are lost relatively fast. On the 100 *τ* time scale, the C–terminal *α*4–helix (residues 69–82) is completely unzipped and fully translocated through the pore. This event is reflected by the loss of native inter–helical contacts with *α*1 (residues 17–28) and *α*2 (residues 35–45) helices and a large decrease of the native contacts of the latter, too. Concomitant with translocation of the *α*4–helix, the *α*3–helix (residues 56–67) is pulled through the contour of the sliding and tightening knotted region, and is thus forced to establish non–native contacts with *α*2 and *α*1 helices. At *t* ≃ 200 *τ*, the *β*2 strand (residues 49–54) begins to slide through the knot, which results in loss of its remaining native contacts and in large formation of non–native contacts. Finally, at *t* ≃ 400 *τ* the knot tightens to as few as ≃ 19 residues and the *β*1 strand (residues 7–12) eventually slides through the knotted region, too. In the low energy barrier pathway, the trefoil knot slides and drops off the chain, which becomes completely untied, at about *t* ≃ 700 *τ*. At this stage about all non–native contacts are removed.

**Figure 3:**
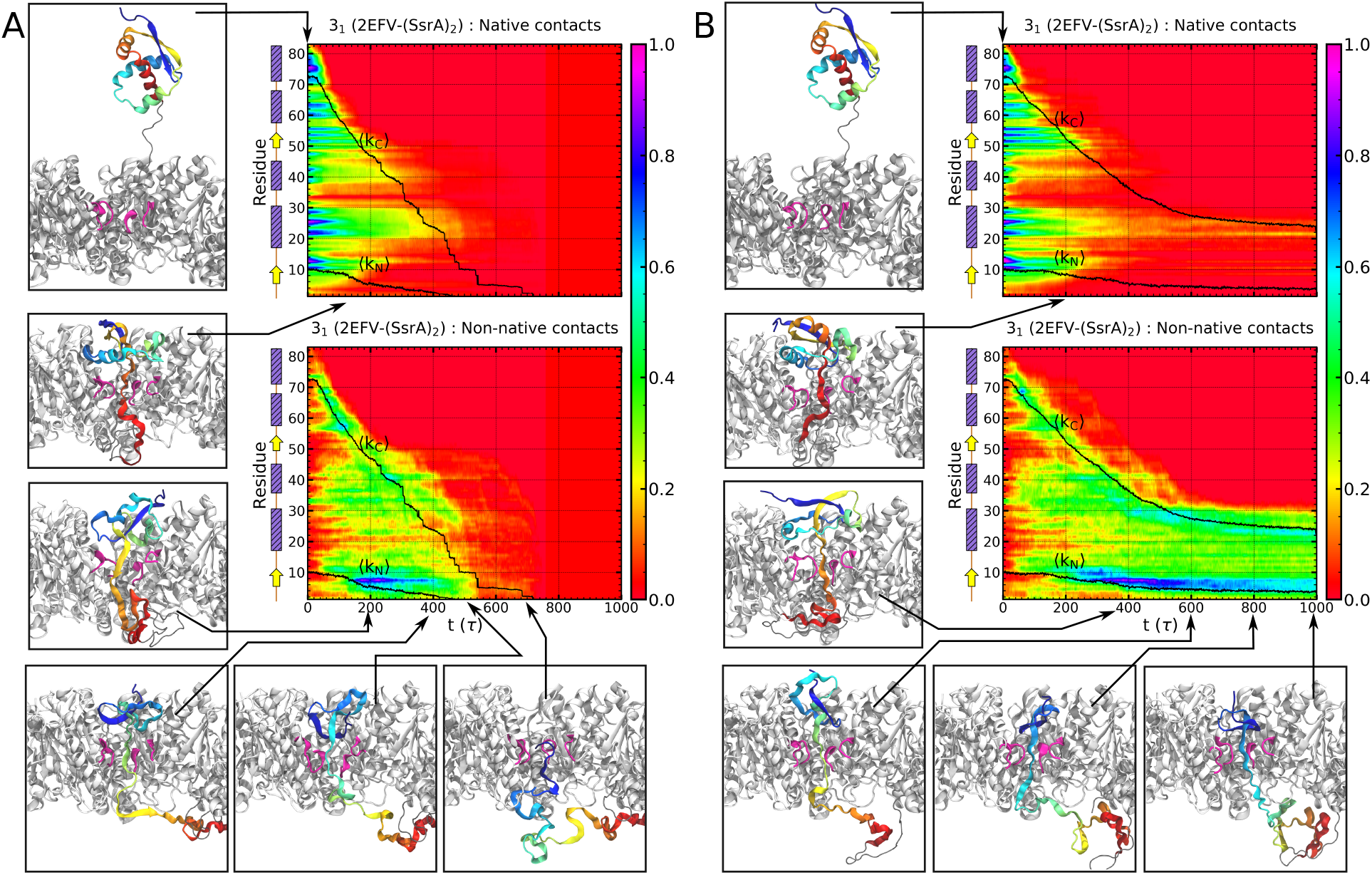
Time evolution of the fraction of native contacts Q_N_ and non–native contacts f_NN_ formed by secondary structural elements of the 3_1_–knotted SP. Ensemble average results are shown for (A) trajectories that result in knot slipping off and (B) trajectories that do not result in knot slipping off. The time–dependent average knot boundaries are indicated using black curves. Representative configurations of the SP (color gradient from red, N–terminal, to blue C–terminal) during unfolding and translocation through the ClpY∆I (grey; central channel loops–purple) pore are also shown.

Our results for the low energy barrier pathway are consistent with recent experimental results, which show that ClpXP can degrade both shallow and deep 3_1_–knotted proteins.^8,9^

In the high energy barrier pathway, native contacts are removed on longer time scales and non–native contacts are present for the entire duration of the simulation. The longest–lived native contacts, which involve residues of *β*1 and *β*2 strands, persist until *t* ≃ 300 *τ* (Figure 3B). After this time, almost no native contacts are present and non–native contacts dominate. By *t* ≃ 650 *τ* the knot slides to the N–terminal and ensnares residues 1–20 through formation of non–native contacts. These strong non–native interactions are not removed on the time scale of our simulations, resulting in stalled translocation in this pathway.

The knot’s sliding dynamics along the SP contour and its stabilization near the N–terminal can be tracked and analyzed through the time evolution of the contact maps. As shown in Figure 4, the native structure is primarily stabilized by the *β*-sheet contacts formed by residues 7–12 and 49–54 of the two strands and inter–helical contacts between the C–terminal *α*4 helix with helices *α*1–*α*3. In accord with the results presented above, the time evolution of these contacts shows that few native contacts survive after *t* ≃ 200 *τ*. In both pathways, native inter–helical contacts are rapidly lost as the *α*4 helix is unzipped and translocated, whereas *β*1–*β*2 contacts persist until this time. We note that, in the higher energy barrier pathway, most native *β*1–*β*2 contacts are still present at this time, whereas, in the lower energy barrier pathway, more non–native contacts are formed by residues of *β*1 and *α*1–*α*2 regions. At *t* ≃ 400 *τ*, additional non–native contacts are formed between *β*1 and *α*1–*α*2 regions in the low energy barrier pathway and extensive non–native contacts are established between *α*1–*α*2 residues, as well as between *β*1 and *α*1–*α*2 residues, in the high energy barrier pathway. After *t* ≃ 600 *τ*, SP unfolding and translocation is completed in the low energy barrier pathway, therefore all contacts are removed, whereas extensive non–native contacts involving *β*1 and *α*2 residues are present in the high energy barrier pathway. A large fraction of these N–terminal non–native contacts persist until *t* ≃ 1000 *τ*, as shown in the final contact map in our trajectories. In particular, formation of salt bridges between Asp17 and Lys20 as well as between Glu4 and Arg7 residues stabilizes the knot and stalls translocation.

**Figure 4:**
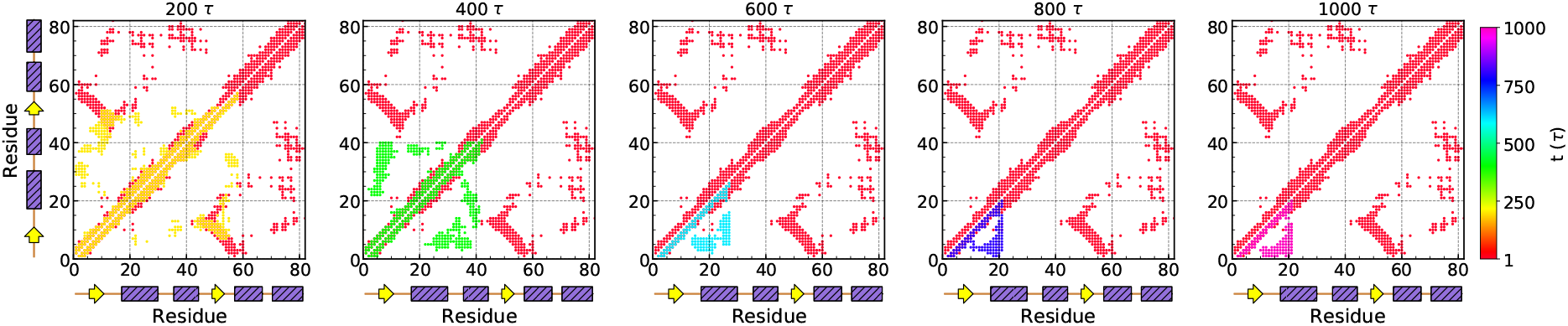
Contact map of the 3_1_–knotted SP in the low and high energy barrier pathways. The time evolution of the contact map of the SP in low (high) energy barrier pathways is shown in the upper (lower) triangle map using time–dependent colors. The native contact map (red) is also indicated in both upper and lower triangles.

These results indicate that strong non–native contacts and self–interactions of the tight knot established along the high energy pathway are the underlying cause for the hindered sliding of the knot itself and of the entire translocation process through the Clp pore.

### Translocation Dynamics of the 3_1_–Knotted SP is Strongly Modulated by both Protein Sequence and Topology

Next, we focus on the time evolution of the translocated chain fraction of the trefoil–knotted SP. To this end, we determine the so–called waiting time, which corresponds to the time spent per monomer in the pore (see Methods). Analyzing the time evolution of this observable allows us to compare and contrast the translocation of knotted SPs to that of homopolymers, which has been extensively characterized.^78–86^ Two dynamic regimes are distinguished in the sequence–wise profile of waiting times. They correspond to an initial (slow) propagation of tension from the monomers inside the pore to the rest of the chain and then to the (fast) translocation of the fully straightened tail. During the first stage, the tension front has to propagate along the “folds” of a self–avoiding polymer,^79,80,87^ which causes a progressive slowing down of translocation or, equivalently, a progressive increase of the waiting times. Translocation instead accelerates during the trail retraction stage, with a progressive decrease of the waiting times.

As shown in the waiting times profiles of Figures 5A–B, translocation of the 3_1_ knotted SP also presents a non–monotonic behavior, with an initial slowing down, followed by acceleration. Despite this qualitative analogy, the differences with the homopolymeric case are major. First, differently from homopolymers, which are defined solely by chain connectivity and excluded volume interactions and can freely fluctuate, SPs have a definite native structure and tension cannot propagate unless native contacts are (continuously) broken in the chain portion that is immediately outside the pore. Concomitantly with the removal of native–contacts, non–native contacts are copiously introduced due to the distortion of the native state and to the sliding and tightening of the knot. These non–native self–interactions, which hinder translocation, reflect in the increasing branch of the waiting time. Consistent with this picture, one notices that the crossover to the accelerating stage occurs in correspondence to the peak of the fraction of non–native contacts, cf. Figs. 2 and 5. After this stage, in fact, the chain is freed from intra–molecular interaction and can translocate fast.

**Figure 5:**
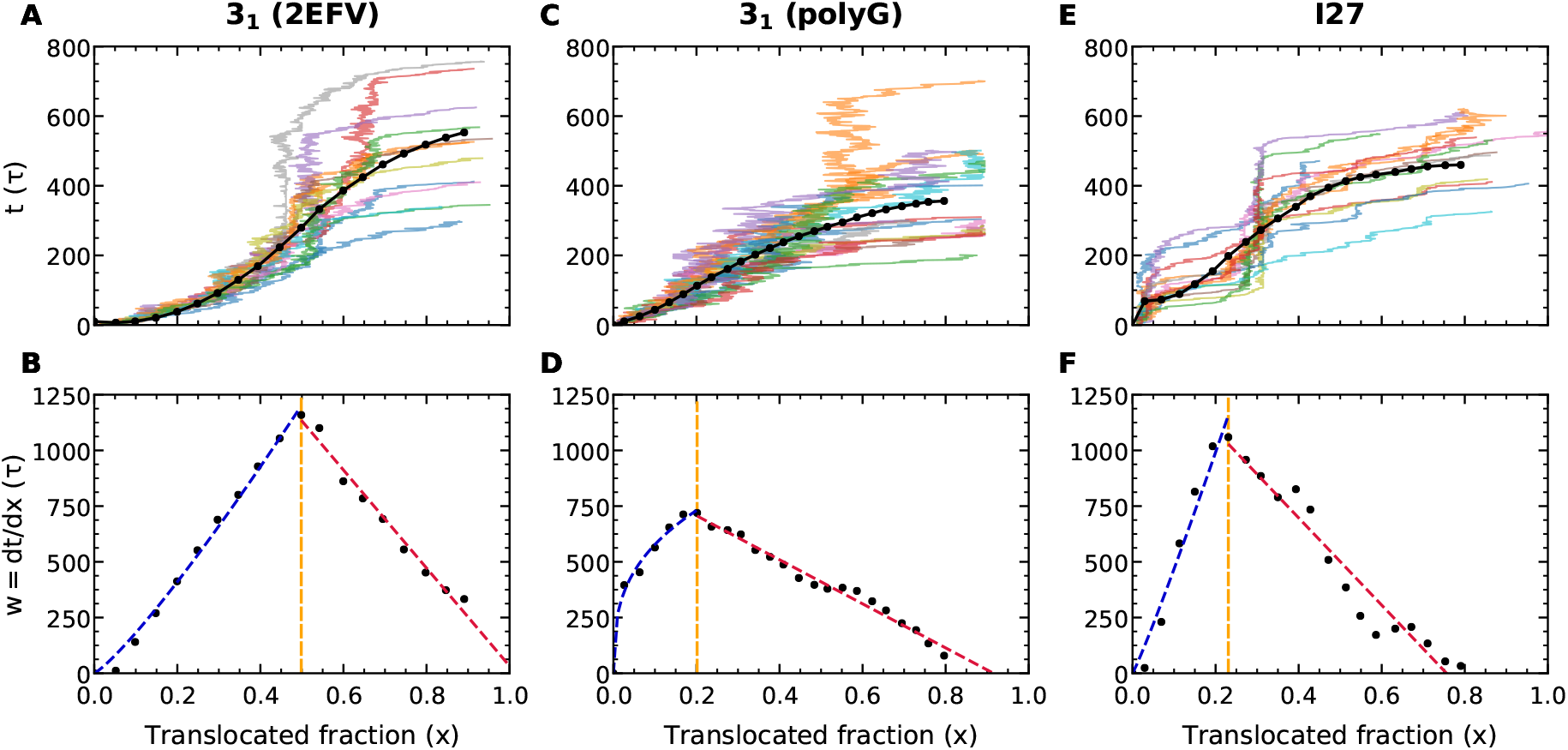
Waiting times of knotted and unknotted proteins. Time evolution and waiting time, *w*, associated with the translocated fraction *x* of the (A)–(B) 3_1_ knotted–SP (C)–(D) 3_1_–knotted polyG and (E)–(F) I27. In the top row, the thin curves indicate individual trajectories and the thick black curves show the averaged values. The bottom plots feature two stages where *w* first increases (blue interpolating line) and then decreases (red line) with time. The two stages are analogous to those observed for homopolymers, where *w* increases during the initial tension propagation along the untranslocated polymer backbone and, once the latter becomes fully rectified, translocation accelerates and *w* decreases.

The structural origin of translocation hindrance is revealed by examining how the waiting times vary with the amino acid index.^88^ As shown in Figure 6A, resistance to translocation in the 3_1_–knotted SP is largely concentrated in the N–terminal region, in accord with our observations in the preceding section of the formation of non–native contacts in this region. Nevertheless, microscopic investigation of the mechanical resistance of the two unfolding pathways reveals quite distinct patterns that involve, on the one hand, a smaller barrier associated with the *β*-sheet that is overcome en route to successful translocation and, on the other, a larger barrier associated with the non–native N–terminal contacts (Figures 6B–C).

**Figure 6:**
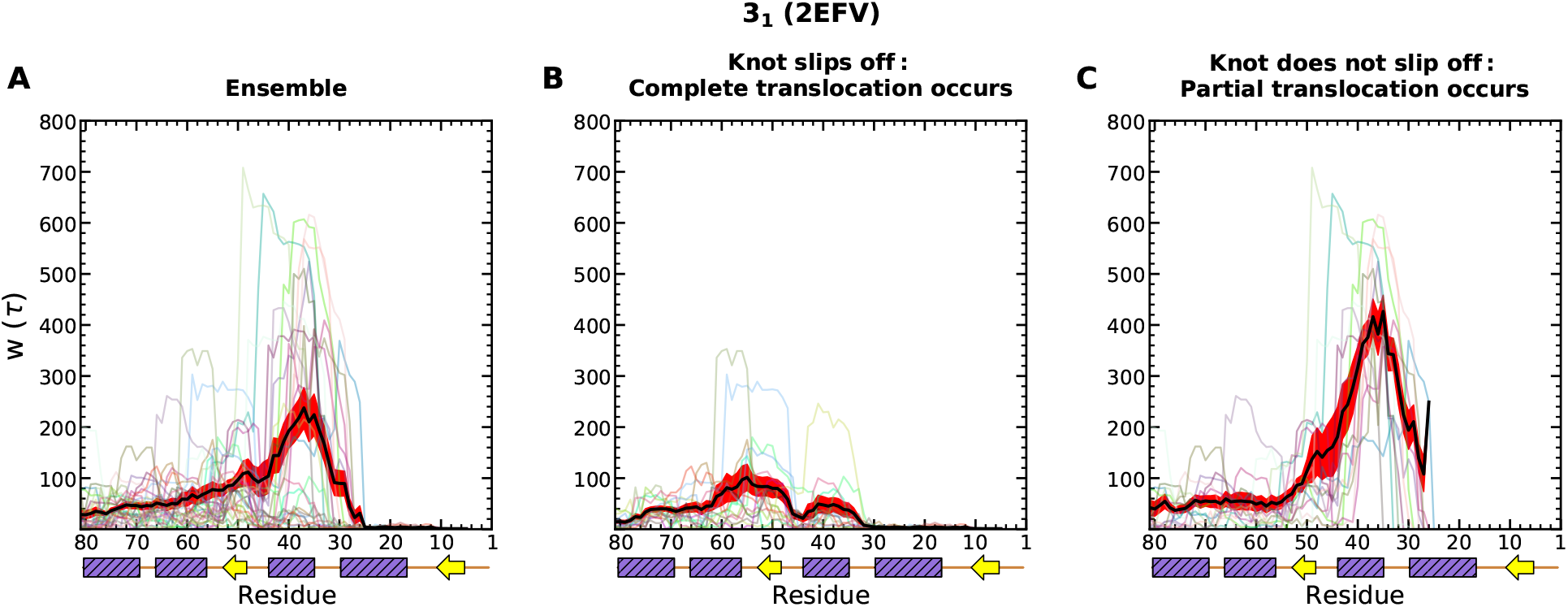
Waiting times per residue in translocation pathways of the 3_1_–knotted SP. The waiting time per residue of the 3_1_ knotted–SP is shown in (A) the ensemble of simulations and in trajectories that result in (B) knot slipping off and complete translocation and (C) knot tightening and partial translocation. Thin curves indicate individual trajectories and the thick black curves show the averaged values. Standard errors are indicated using red bands.

Given the above considerations, it is of interest to establish to what extent the observed hindrance is due to the knotted topology of the protein backbone and to the protein sequence. To isolate and separate the two contributions, we compared the translocation response of MJ0366 with that of a polyG variant of the same trefoil–knotted SP and then of an I27 domain. The polyG variant was obtained by removing “in silico” the side chain of the MJ0366 while the I27 domain, often considered in single–molecule experiments, is globular and unknotted (see Methods and Table 1). Used as terms of comparison, these two entries can thus highlight sequence– and topology–dependent effects.

A first set of comparative results are presented in Figure 5. Inspection of the translocation curves in Figures 5C,E show large dispersion of the trajectories for I27, associated with cooperative removal of inter–strand contacts in this unknotted entry. Most of all, one observes that without the trefoil knot (I27) or without side chains (polyG), the waiting time profile is modified in much the same way. The location of the waiting time peak, which occurs for *x* ∼ 0.5 for the trefoil–knotted SP, is shifted to the much smaller *x* ∼ 0.2 for both I27 and polyG. The lower *x* values of the peak location (and of the peak height too for polyG) indicate that translocation is facilitated in these cases.

As shown in Figure S2, mechanical fingerprints of the polyG and I27 vividly reflect the response of these SPs to the axial pulling forces. Whereas little resistance can be offered by the polyG structure, which is less prone to form non–native contacts, resistance to translocation in the I27 features prominently the stringent role of inter–strand contacts.

We thus conclude that the hindrance observed in the translocation of the knotted SP is not due to backbone topology alone (because the polyG response is different, as also indicated in Figures S3 and S4 and Movie S3), nor to general side chain interactions alone (because the I27 response is different too). The above comparison instead points at a synergistic contribution of the knotted backbone topology *and* the side chain interactions. This previously–unknown effect, which we can highlight thanks to the atomistic detail of the model, can be rationalised by considering that it is not only the mere tightening of the knot that slows down translocation, but that the dominant hindrance (reflected in the shift of both height and location of the waiting time peak) is introduced by the side chains being dragged along the tightly knotted chain contour.

### Knot Tightening Drives Dynamic Coupling of Translocation and Tension Propagation

Microscopic events that underlie the distinct translocation behavior of the compared entries are aptly illustrated by the time evolution of the orientation of virtual C_*α*_–C_*α*_ bonds with respect to the axial force direction.^89^ As shown in Figures 7A–C, for globular SPs, Clp–mediated pulling elicits strong mechanical response throughout the protein structure which is reflected in the extensive alignment of virtual bonds with the direction of pore axis. Mechanical pulling applied at the C–terminal of these SPs results in rapid alignment of *β*–strand G of I27 and *α*4 helix of the 3_1_–knotted protein with the direction of the external force. Along with these regions, which directly experience the ClpY loop–mediated force, the *β*1–strand and *α*1 − *α*3 helices in the 3_1_ SP and the *β*–sheet that includes the I27 terminals also become aligned with the force direction.

**Figure 7:**
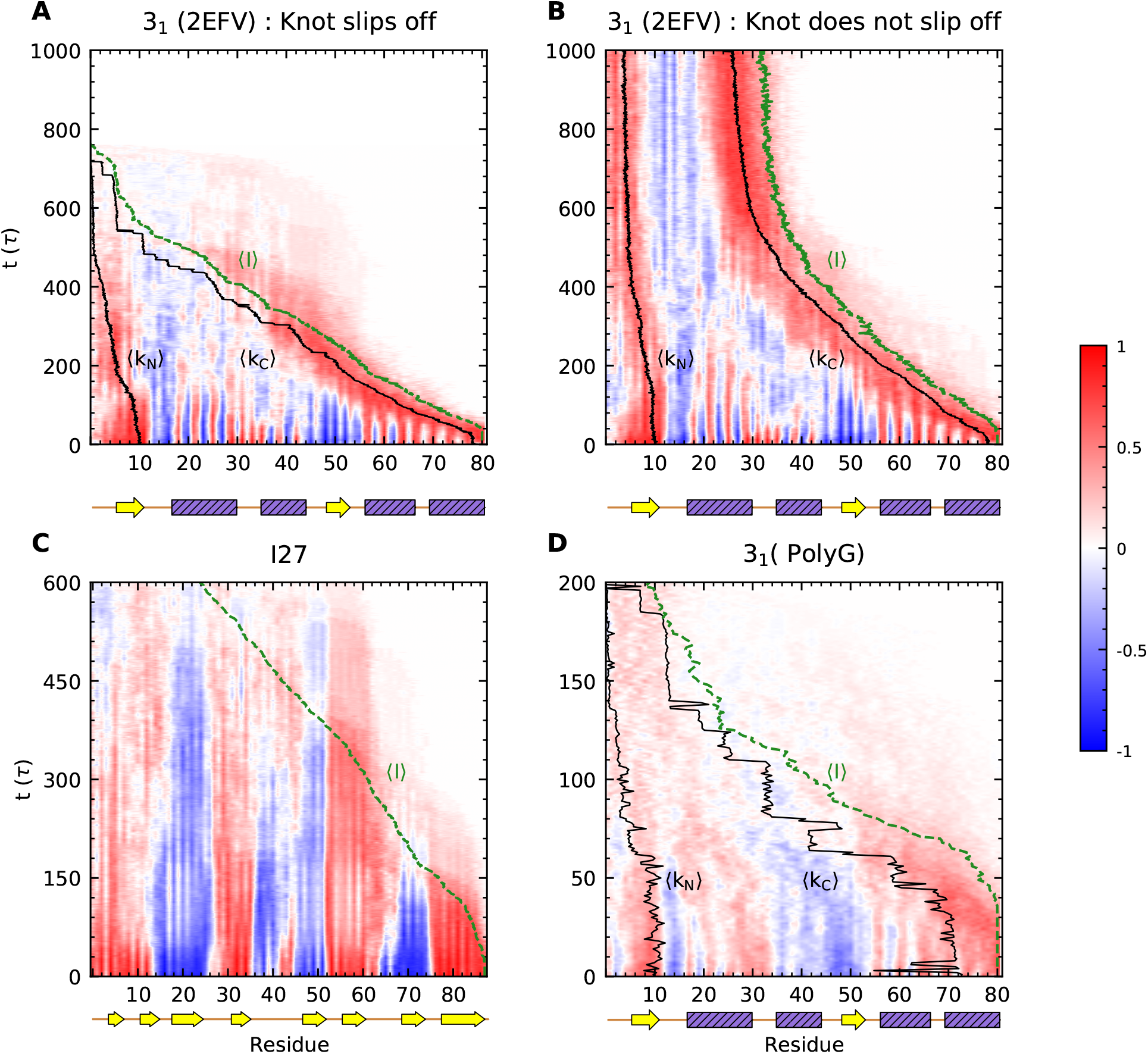
Tension propagation within the 3_1_–knotted SPs and I27. Average time–dependent orientation of virtual C_*α*_–C_*α*_ bonds involving bonded amino acids, cos *θ*, is indicated with respect to the axial force direction in trajectories that (A) result or (B) do not result in knot slipping off the chain. The average knot boundaries (solid lines, black) and the average translocation line (dashed line, green) are also indicated. (C)–(D) Same as in (A)–(B) for (C) I27 and (D) polyG substrates. Secondary structural elements are indicated for each protein (for polyG, the 3_1_ structure is indicated as a reference).

As translocation proceeds, the C–terminal knot edge, delineated by the 〈k_C_〉 sequence position, is firmly engaged by the ATPase and primed for translocation by the short polypeptide segment, upper bounded at position 〈*I*〉, that spans the length of the pore and emerges on the *trans* pore side (see Methods). Dynamically, more core residues in each SP are forced to align while approaching the pore until the fully or partially tightened knot slips off the chain or stalls translocation (Figures 7A–B) or the I27 folded domain unravels completely (Figure 7C). Notably, in the case of the knotted SP, it is not only the region closest to the pore (and at the proximal knot edge, k_C_) that re-orients and aligns with the pore axis, but the same occurs at the distal knot edge, k_N_, too. The effect is stronger for the trefoil–knotted MJ0366 than its polyG variant (Figures 7A–B,D). This divergent behavior is attributed to the weaker engagement of the polyG SP, indicated by the larger distance along the polypeptide chain between the translocation boundary, 〈*I*〉, and the proximal knot edge, k_C_. Alignment at the free N–terminal is also observed in I27, however, in this case, it is directly related to strong hydrogen bond contacts formed between A′–G strands that connect the protein ends (Figure 7C). Loss of the stabilizing N–C interface and translocation of the C–terminal strand G, which occur on a time scale of ≃ 300 *τ*, significantly weaken the mechanical response of most structural regions as indicated by softer alignment of individual virtual bonds (Figure 7C).

The results suggest that the wiring of the knotted backbone can propagate mechanical pulling to the knot edge away from the pore while, at the same time, the chain portion between the two edges is mostly undisturbed.

### Stabilization of Non–Native Structure and Complex Topology of 5_2_ and 6_1_ Knots Hinder Long–Distance Tension Propagation

We next consider two proteins with higher topological complexity than the trefoil knot, namely the 5_2_–knotted ubiquitin carboxy–terminal hydrolase L1 (UCH–L1) and the 6_1_–knotted *α*–haloacid dehalogenase I SPs (Figure 1). For both cases we found necessary to step up the average magnitude of pulling forces to 500 pN, since forces of 300 pN are not conducive to unfolding and translocation over the simulated time scales (see Methods, Table 1 and Supporting Information Movies S4–S5). For the UCH–L1 SP we consider Clp–mediated processing in both N–C and C–N directions, as probed in experimental studies.^9–11^

The native structure of the 5_2_–knotted UCH–L1 comprises seven *α*–helices wrapped around a six–stranded *β*–sheet core. The repetitive ClpY–mediated pulling at the N–terminal causes the unfolding and translocation of helices *α*1 and *α*2 and of strands *β*1 and *β*2, with a resulting destabilization of the native structure (see also Supporting Information Movie S4). This is reflected by the large loss of native contacts shown in Figure 8, with only about a third of them, 〈Q_N_〉 ≃ 0.3, surviving on the simulated time scale, *t* = 1000 *τ*. Notice that the loss of native contacts occurs in spite of a limited progress of translocation, the translocated chain fraction being only about *x* ∼ 0.3 at *t* = 1000 *τ*. The process is presumably hindered by the numerous non–native contacts, 〈f_NN_〉 ≃ 0.5, that replace the lost native ones. As in the case of the trefoil–knotted SP, most non–native contacts involve side chain interactions, which highlights their consistent involvement in hindering translocation and knot sliding.

**Figure 8:**
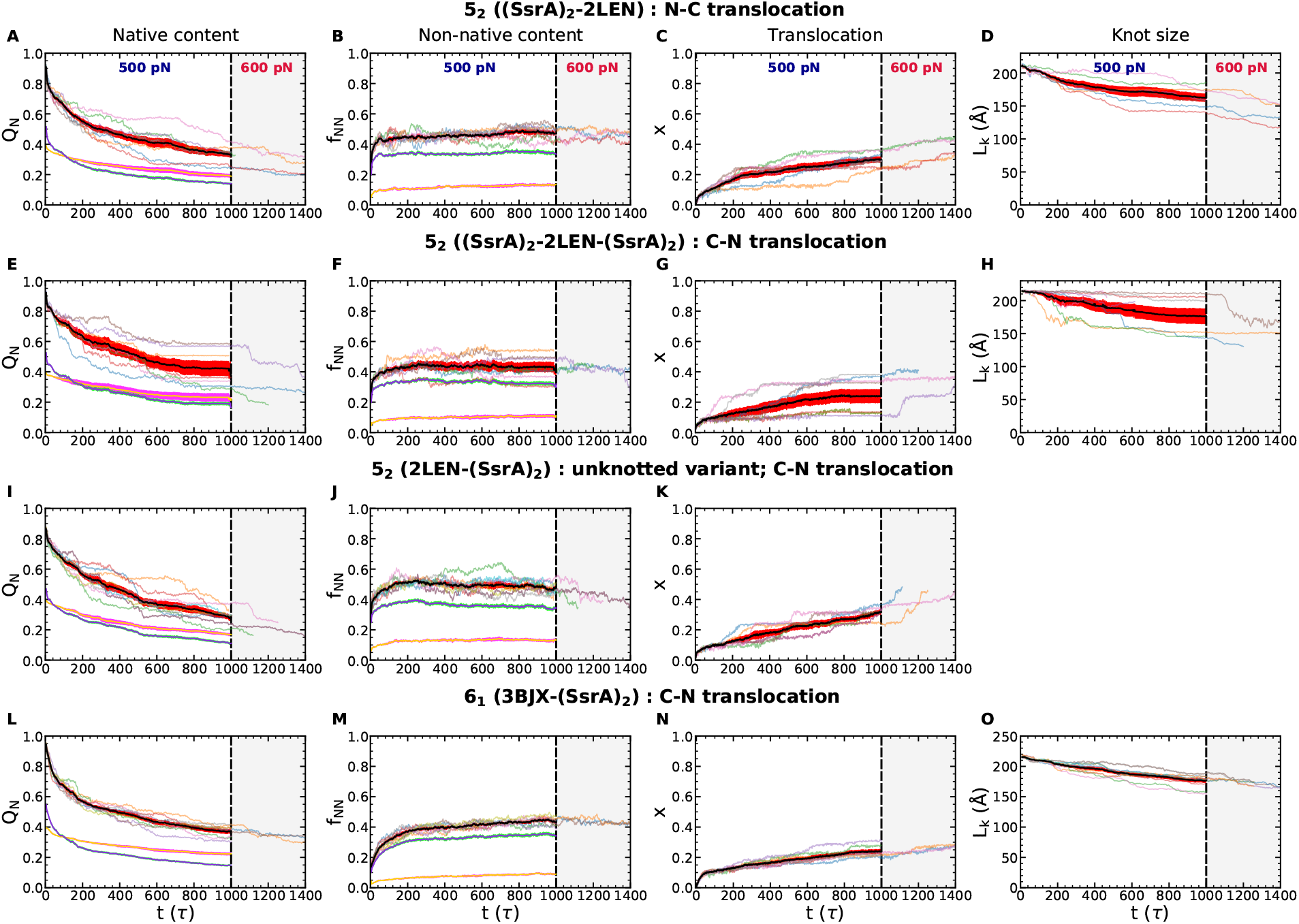
Unfolding and knot tightening in 5_2_– and 6_1_–knotted SPs. Time evolution of (A) the fraction of native contacts Q_N_ (B) the fraction of non–native contacts f_NN_, (C) translocated fraction *x* and (D) the knot length L_K_ for the 5_2_–knotted SP processed in the N–C direction. (E)–(O) Same as in (A)–(D) for (E)–(H) the 5_2_–knotted SP processed in the C–N direction; (I)–(K) the SP variant with untied 5_2_–knot processed in the C–N direction; (L)–(O)) the 6_1_–knotted SP processed in the C–N direction. Simulations include repetitive axial pulling forces with average magnitude of 500 pN for *t* < 1000*τ* (see Methods). Fast–forwarding of selected simulation trajectories was performed, for 1000 *τ* < *t* < 1400 *τ* (shaded region), by using axial pulling forces with average magnitude of 600 pN. Individual traces (thin curves) and averages and standard errors (thick curves; black and red, respectively) are shown. Average contribution (and standard error) of contacts involving backbone–backbone interactions (yellow and red) and side chain–side chain or side chain–backbone interactions (purple and cyan) are also indicated.

At *t* = 1000 *τ*, the average knot size has reduced to 〈L_K,min_〉 ≃ 167.7 ± 3.9 Å, a 10% larger value than the L_K_ ≃ 146 Å knot size observed in experiments where UCH–L1 was mechanically stretched from both ends using optical tweezers.^48^ As shown in Figure 9A and Supporting Information Movie S4, in Clp–mediated pulling the knot size is reduced by approximately 50 amino acids compared to the initial state. One notes that the tightening of the knot is induced by the sliding of only one of its boundaries, namely the one proximal to the pore, 〈k_N_〉, whereas the distal C–terminal end, 〈k_C_〉, remains nearly stationary. In fact, the 〈k_N_〉 boundary always remains at a short sequence distance from the protein region just exiting the pore, 〈*I*〉. Thus, the larger knot size observed here compared to stretching experiments reflects the asymmetric pulling of SPs’ N– and C– regions during translocation. Notably, almost all regions of the knotted domain experience re–orientations as the folded structure deforms (Figure 9A).

**Figure 9:**
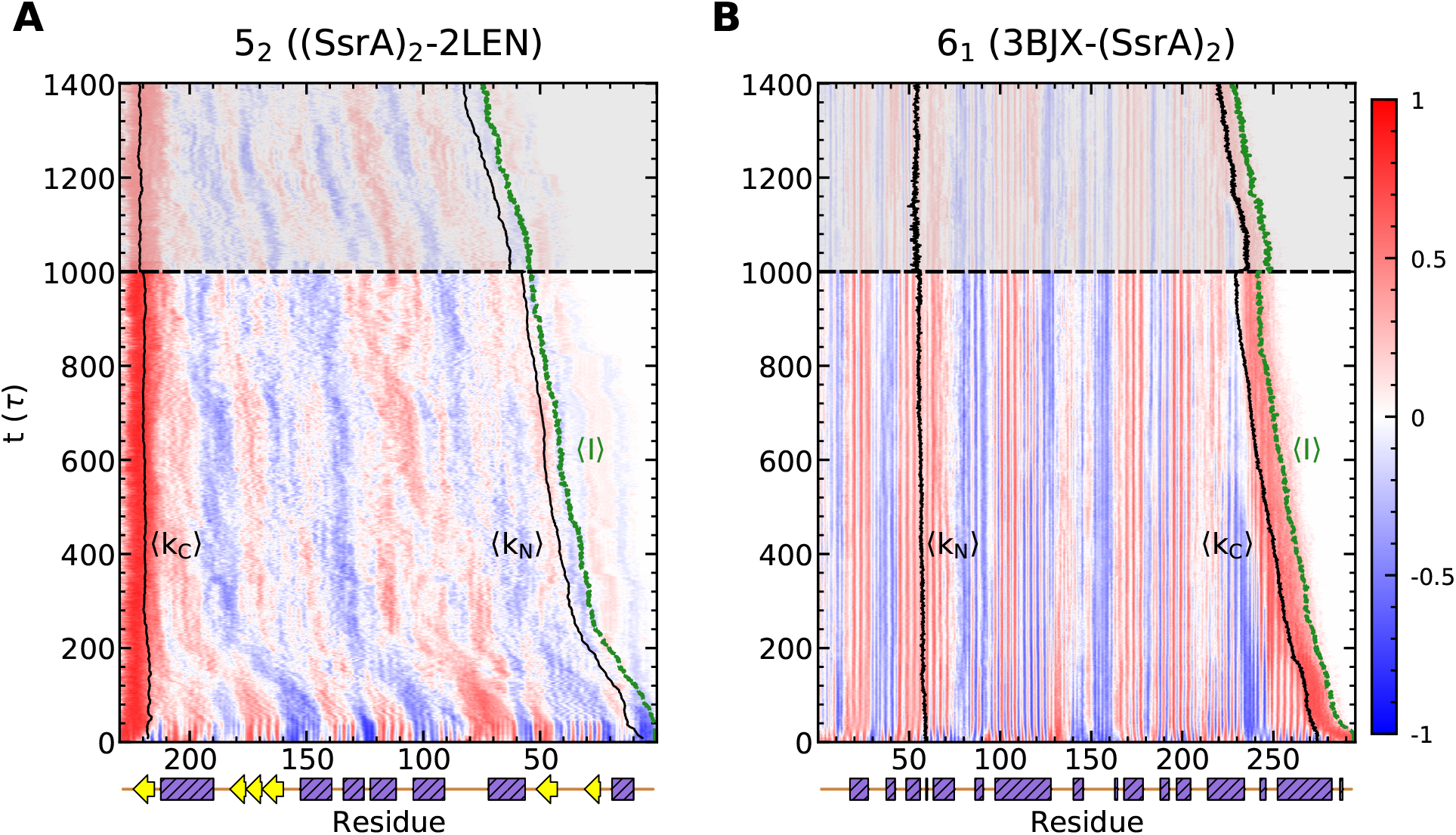
Tension propagation within the 5_2_ and 6_1_–knotted SPs. Average time–dependent orientation of virtual C_*α*_–C_*α*_ bonds involving successive amino acids, cos *θ*, is indicated with respect to the axial force direction for (A) the 5_2_–knotted SP processed in the N–C direction and (B) the 6_1_–knotted SP processed in the C–N direction. The average knot boundaries (solid lines, black) and translocation line (dashed line, green) are also indicated. The shaded region corresponds to fast–forwarding of selected trajectories by using larger axial pulling forces.

As shown in Figure 10A, there is only limited coupling of the structural deformations in the central protein region (residues 50–100) and the C–terminal one (residues 150–231). Whereas the former loses a large fraction of native contacts, the latter mostly retains its native content, except near the 〈k_C_〉 edge. Even before the disruption of the native contacts, the sliding and tightening of the knotted region establishes several long–lived non–native ones that arguably contribute to the slow progress of translocation along with the more complex and larger structure of UCH–L1.

**Figure 10:**
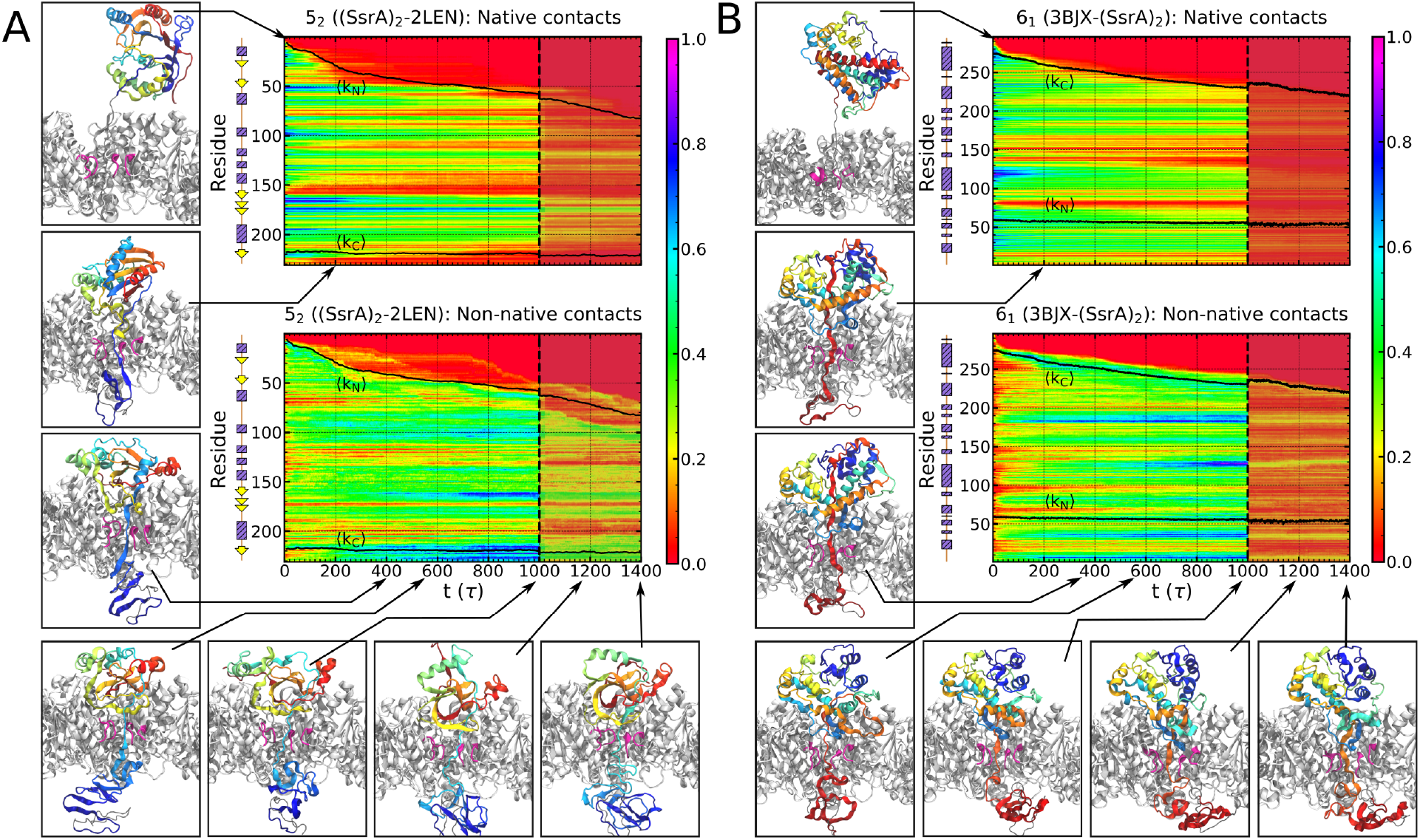
Time evolution of the fraction of native contacts (Q_N_) and of the fraction of non–native contacts (f_NN_) for the mechanisms of unfolding and translocation of (A) the 5_2_–knotted SP in the N–C direction and (B) the 6_1_–knotted SPs in the C–N direction mediated by ClpY∆I. Also shown are representative configurations of the ClpY∆I–SP system during simulation trajectories. The shaded region corresponds to fast–forwarding of selected trajectories by using larger axial pulling forces.

To enhance sampling of unfolding and translocation events, we extended several simulation trajectories from *t* = 1000 *τ* up to *t* = 1400 *τ* while using pulling forces with the larger average magnitude of 600 pN. As shown in Figure 8, this fast–forwarding approach yields an appreciable advancement of translocation, ∆*x* ≃ 0.1, concomitant with a reduction of the native contacts, ∆*Q*_*N*_ ≃ 0.1. The larger pulling force also enhances knot tightening, ∆*L*_*K*_ ≃ −25Å, analogously to from what was observed in force ramping experiments using optical tweezers setups.^48^ The fact that both the knot sizes and the fraction of non–native contacts eventually reach a plateau suggests that completing the passage of the polypeptide chain through the pore requires coordinated translocation steps, as noted in the case of the 3_1_ knot.

We also probed Clp–mediated processing of the 5_2_–knotted SP in the C–N direction. In this setup, the distal edge of the knot, k_N_, is located only a few amino acids away from the free (unpinned) N–terminal, and thus the shallow knot can become spontaneously unknotted due to thermal fluctuations. We took advantage of this unknotting propensity of UCH–L1 knot to compare and contrast ClpY–mediated unfolding and translocation of two variants of the SP: one without the shallow knot (obtained from runs where the SP spontaneously untied) and one where the 5_2_ knot was made deeper (and thus preserved) with an auxiliary segment attached at the distal end (see Methods and Table 1). The two variants have thus very high structural similarity but distinct topology.

The C–N unfolding and translocation behavior of the knotted and unknotted UCH–L1 variants are shown in Figures 8E–F and 8I–J and Movies S6–S7. One observes that the knotted SP variant retains a larger fraction of native contacts, 〈Q_N_〉 ≃ 0.45, and has fewer non–native ones, 〈f_NN_〉 ≃ 0.45 than the unknotted variant, for which 〈Q_N_〉 ≃ 0.35 and 〈f_NN_〉 ≃ 0.5.

Because of the resistance offered by the knot, about 50% (4) of the trajectories show negligible translocation, *x* ≲ 0.05 at the end of the trajectories, and the minimum knot length is 〈L_K,min_〉 ≃ 169.6 ± 14.1 Å, similar to the N–C directed translocation (Figures 8G–H and 8K). Correspondingly, a larger translocated fraction is observed for the unknotted variant.

Although the absence of the knot is expected to reduce the SP’s sliding friction, we note that, by itself, it is still insufficient to permit complete SP unfolding and translocation over the simulated timespan. In fact, the C–N translocation evolution is very similar for the unknotted and unknotted variants of UCH–L1. The fact that the sliding kinetics of the unknotted variant is not faster, indicates that translocation is not necessarily hindered by complex topologies *per se*, but rather by more general and pervasive forms of entanglement that typically, but not exclusively, accompany proper knots. For UCH–L1, which is tied in a twist knot topology, these forms of entanglement include the essential (twist) crossings of the polypeptide chain (Figure 1B) which persist after the knot–untying strand passage at the C–terminal, and whose inherent friction can hinder the driven unfolding process. The result bears qualitative analogies with the case of xrRNAs, a class of exonuclease resistant viral RNAs^90^ that can resist translocation at the 5′ end by virtue of their complex architecture and network of intramolecular interactions. ^88,91^

As shown in Figures S5A–B, secondary structural elements retain most of their native contacts and form relatively fewer non–native contacts in the knotted SP variant, which, strikingly, sustains very modest chain re–orientations, too. The latter fact indicates weak tension propagation, and is in accord with the limited translocation observed in this case (Figure S6A). By contrast, discernible changes in chain orientation occur for the portions of the unknotted variant upon approaching the pore (Figure S6B).

Figures 11A–B show that most maxima of the waiting times profiles occur in correspondence or in the vicinity of *β*–strands. This occurs for both N–C and C–N pore entries, indicating that the rupture of these elements poses a significant obstacle to the translocation process regardless of the process directionality. However, the location of the peaks depends strongly on the entry direction. Note, in particular, that the peaks’ location for C–N translocations of the knotted and unknotted version of UCH–L1 are very similar (Figure 11B–C). The analysis thus exposes the role that specific secondary structural elements have in the translocation process in addition to the backbone topology. This mechanical fingerprint, resolved in microscopic detail, is also valuable to rationalize the directional response observed in the aforementioned experiments on 5_2_–knotted SPs^9–12^ and clarify how, beyond topology, it is deeply rooted in the secondary and tertiary organization of the SPs which typically present a completely different succession of motifs from the two ends.

**Figure 11:**
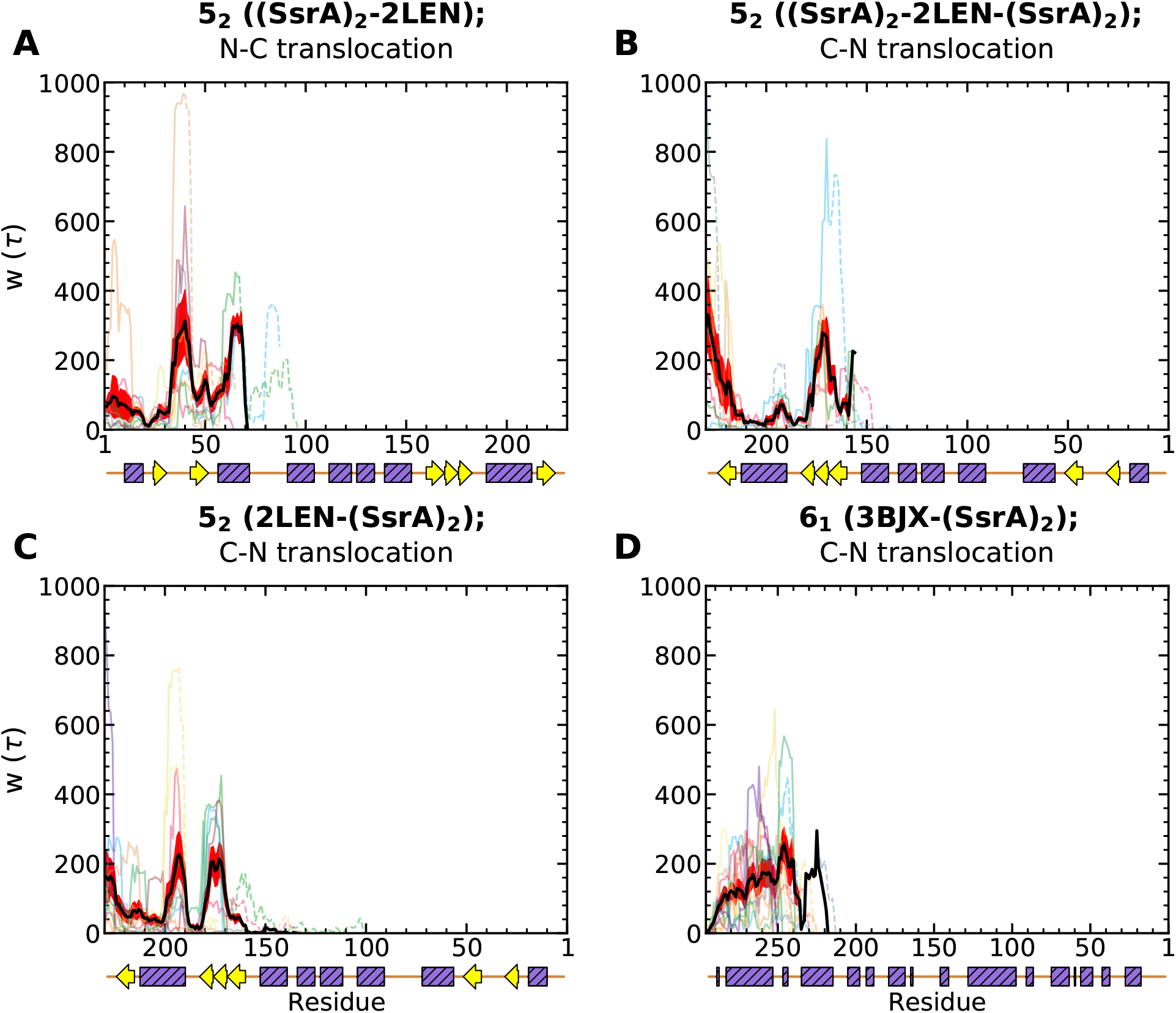
Translocation hindrance in directional processing of the 5_2_ and 6_1_–knotted SPs. Time series of the waiting times per residue for (A)–(B) the 5_2_–knotted SP processed in (A) the N–C and (B) the C–N direction, (C) the unknotted variant of the 5_2_ SP processed in the C–N direction, and (D) the 6_1_–knotted SP processed in the C–N direction. The waiting times (thin curves) for individual trajectories are shown using solid lines, for *t* < 1000*τ*, and dashed lines, for 1000 *τ* < *t* < 1400 *τ*. The average waiting time (black curve) and standard error (red) are also indicated for *t* < 1000 *τ*.

As in the case of the 5_2_–knotted UCH–L1, unfolding of the 6_1_–knotted SP results in large loss of native contacts, replaced by persistent non–native ones, though their incidence is smaller than for UCH–L1, 〈f_NN_〉 ≃ 0.45 (Figure 8). Partial translocation of this predominantly *α*–helical domain is limited to the two C–terminal helices, yielding an average translocated fraction of 〈*x*〉 ≃ 0.2 and average knot length of 〈L_K,min_〉 ≃ 173.3 ± 2.4 Å. After translocation of the two helices, the translocation process becomes apparently stalled, and the k_C_ edge of the knot becomes tightly entangled (Supporting Information Movie S8).

Fast–forwarding of simulation trajectories, using axial pulling forces with average magnitude of 600 pN, indicate additional translocation upon further knot tightening. While this behavior is in accord with observations for the 5_2_–knotted SPs, we note that, in the case of the 6_1_–knotted SP, translocation hindrance emerges from non–specific entanglement of helical structure (Figure 11D).

Notably, SP unfolding is mostly localized at the C–terminal region, whereas the untranslocated SP fragment largely maintains its shape and orientation. Thus, in contrast to the UCH–L1 SP, the direction of the applied force remains nearly constant throughout the folded region except for the regions that approach and eventually enter the pore (Figures 9B and 10B). Mechanisms involving the 5_2_– and 6_1_–knotted SPs highlight the relationship between translocation and knot complexity and are in accord with the trends noted above for the translocated fraction of the 3_1_–knotted SP. Consistently, as shown in Figure 9, the limited translocation of the 5_2_– and 6_1_–knotted SPs results in time–dependent changes in orientation only near the knot edge actively engaged by the Clp ATPase. This pattern of weak transmission of tension to the distal end of the knot is in accord with the systematic studies of translocation of polymers with 5_2_ and 6_1_ knots,^54^ which indicate that the pulling force is predominantly experienced near the pulled terminal with only a modest transmission of tension along the knotted backbone and the network of contacts.

## Discussion and Conclusions

In this computational study we addressed, with atomistic detail, key aspects of the degradation of entangled proteins effected by Clp biological nanomachines. We focused on how knotted proteins are translocated through the ClpY lumen in repetitive allosteric cycles of the degrading complex and addressed the following questions: What is the relative contribution of the protein’s structure (knotted backbone, secondary structure elements) and sequence (side chains’ size and interactions) on the translocation compliance of knotted proteins? What role do non–native interactions play in the process? How likely is it that tight protein knots can slip through the flexible ClpY pore?

To define and address these questions, we built on the numerous studies that, using coarse–grained models, have established the general physical underpinnings of how knotted filaments respond when pulled through narrow rigid pores. The chain tension propagating from the pore, and the steric interaction between the chain and the pore, both cause the tightening of the knot at the pore entrance. For translocation to proceed, the substrate must slide along its knotted contour and overcome the multiple barriers associated with the tightly contacting regions inside the knot. ^50–52,56^ Depending on the magnitude of the driving forces^51^ and knot complexity,^54^ the resulting topological friction can be large enough to stall translocation.

Here, we leveraged atomistic simulations to reveal novel aspects of the translocation of knotted proteins through biological nanomachines such as ClpY. We specifically focused on how translocation compliance is affected by: (i) the role of the backbone architecture and topology versus that of the side chains, (ii) the role of non–native interactions and (iii) the plasticity of the ClpY lumen.

For our assessment we relied on several comparative analyses. To ascertain the effects of entanglement complexity, we considered representatives of three different types of backbone topologies: the trefoil (3_1_), three–twist (5_2_), and stevedore (6_1_) knots, as well as unknotted SPs. The knotted entries corresponded, respectively, to MJ0366, UCH–L1 and *α*—haloacid dehalogenase I from *Pseudomonas putida*. The unknotted entries included the I27 globular domain, which is similar in size to the trefoil–knotted MJ0366 entry but biologically unrelated to it, and a conformer from the molecular dynamics evolution of the 5_2_-knotted UCH-L1 where the shallow knot had spontaneously slipped off the N terminal. Finally, to establish the relative role of backbone and side chains, we contrasted the behavior of the trefoil–knotted protein with a polyG version of it obtained by shaving off *in silico* all sidechains of MJ0366, thus preserving the native backbone geometry and 3_1_ topology.

Our results highlight strong translocation modulation by intramolecular interactions, especially those involving side chains, and plasticity of biological pore. Comparative analysis of the behavior of the native trefoil–knotted protein, of the polyG variant, and of the unknotted I27 domain, show that side chain interactions are as important as the backbone knotted topology in determining the translocation compliance. Strikingly, the synergistic action of *both* side chain interactions and the backbone knotted state can hinder translocation to a much greater extent than either of the two terms.

In addition, we find that interactions involving side chains can establish long–lived non–native contacts that can trap the unfolding and translocation process of the substrate protein. For instance, non–native interactions give rise to bifurcation of unfolding and translocation pathways of the trefoil–knotted SP into faster or slower processes and stalled translocation of the more complex 5_2_– and 6_1_–knotted SPs.

We also found that dynamic flexibility of the biological pore, controlled by allosteric cycles of the nanomachine, enables re–orientation of the SP and application of the pulling force along directions of low mechanical resistance. For the 5_2_–knotted UCH–L1 the re–orientation compliance depended strongly on the pulling direction: it was negligible for C to N translocation and very conspicuous for the N to C one. Though for this topology, only a partial translocation could be obtained, the results are consistent with the experimentally–observed higher UCH–L1 susceptibility to unfolding and degradation from the N end.^9^

We noted that native and non–native intramolecular contacts prevent the effective propagation of tension along the chain as it normally happens in fluctuating homopolymers. We do observe, however, that complex backbone architectures allow for the “leapfrog” transmission of tension to parts of the chain that are distant from the pore.

Despite the intrinsic flexibility of the ClpY lumen, we did not observe translocation of the entire knotted portion in our simulations, not even for the smallest knotted SP considered. Though it is possible that complete knot translocations can occur, it is unlikely that these events are part of the dominant degradation pathways. The behavior of UCH–L1, both its knotted and unknotted variants, indicates that their topological and geometrical complexities are very effective in slowing down unfolding and translocation progress. We cannot rule out that, over sufficiently–long and currently inaccessible time scales, the degradation progress can become stalled. In such drastic circumstances, it is still possible that the ATPase–operated cleaving of the SP portion that successfully translocated, could still lead to the spontaneous untying of the entanglement of the undegraded SP remainder, once the latter is released. ^28^ This mechanism is highly plausible given the propensity of proteases or the proteasome to release partially degraded fragments of substrates that require excessive processing.^92–97^ We thus speculate that the same mechanism could lead to the spontaneous simplification of the (shallower) entanglement of the released undegraded substrate, which could then be more easily and fully processed by proteases upon recapture.

More generally, our atomistic study indicates that backbone geometry, side–chain interactions, non–native contacts and translocation directionality, can all contribute to impair ATPase–effected degradation alongside with the SP knotted topology, which is thus neither the sole nor an indispensable element for hindering translocation. The observation poses two natural questions, namely: what other forms of entanglement, beyond knots, can stall or thwart the translocation process of naturally–occurring proteins? Is it at all possible to recapitulate the observed sequence–specific contribution to translocation hindrance (e.g., side–chain interactions and non–native contacts) with coarse–grained models, which would allow reaching much longer time scales? Both questions touch central aspects of proteins’ functionality and mechanics and we believe would be ideal avenues for future studies.

Further extensions of our study could also involve an increased complexity of the ClpY cycle description by adding ATP–independent mechanisms, such as Brownian ratchets,^98,99^ by removing the constraint of strict processivity, or by including transitions to and from intermediate states whose structure might be solved and become available in the future. The high resolution afforded by our simulations can provide particularly useful roadmaps for future experimental studies probing these questions. Specifically, single–molecule experiments may afford direct measurements of the knot length, pauses during translocation, or the length of the translocated polypeptide segment.^48,100–102^ In addition, the detailed steps of the unfolding and translocation reactions could be addressed with experiments that compare processing of wild–type knotted SP and their variants that controllably alter single mechanical interfaces to either block specific unfolding pathways or to weaken critical contacts. ^8,11,48^

## Acknowledgement

G.S. expresses his gratitude for the inspiration that D. Thirumalai provided through his mentorship and passion for the scientific pursuit. We thank Sue Wickner and Mike Maurizi for stimulating discussions. This work has been supported by the National Science Foundation grant MCB–1516918 to G.S. This work used the Extreme Science and Engineering Discovery Environment (XSEDE), which is supported by NSF grant number ACI–1548562. XSEDE Comet resources at the San Diego Supercomputer Center and Bridges resources at the Pittsburgh Supercomputer Center were used through allocation TG–MCB170020 to G.S.

## Supporting Information Available

Sampling of unfolding and translocation pathways in simulations of the 3_1_–knotted SP using axial pulling forces with average magnitude of 200 pN and 300 pN (Figure S1); waiting times per residue for I27 and polyG SP (Figure S2); time–dependent contact map of the 3_1_–knotted polyG SP (Figure S3); time evolution of the fraction of native and non–native contacts formed by secondary structural elements of the 3_1_–knotted polyG SP (Figure S4); time evolution of the fraction of native and non–native contacts formed by secondary structural elements (Figure S5) and tension propagation (Figure S6) in variants of the 5_2_–knotted SP engaged at the C–terminal.

Movie S1. Knot slip–off and complete translocation of the 3_1_–knotted SPs mediated by ClpY∆I

Movie S2. Tightening and partial translocation of the 3_1_–knottted SP mediated by ClpY∆I

Movie S3. Knot slip–off and translocation of the 3_1_–knotted polyG SP mediated by ClpY∆I

Movie S4. Tightening and partial translocation of the 5_2_–knotted SP processed in the N–C direction by ClpY∆I using pulling forces with average magnitude of 500 pN.

Movie S5. Absence of unfolding and translocation of the 5_2_–knotted SP processed in the N–C direction by ClpY∆I using pulling forces with average magnitude of 300 pN.

Movie S6. Unfolding and partial translocation of the unknotted SP 2LEN processed in the C–N direction by ClpY∆I

Movie S7. Tightening and partial translocation of the 5_2_–knotted SP processed in the C–N direction by ClpY∆I

Movie S8. Tightening and partial translocation of the 6_1_–knotted SP processed in the N–C direction by ClpY∆I.

