## Supporting Information for "Unfolding and Translocation of Knotted Proteins by Clp Biological Nanomachines: Synergistic Contribution of Primary Sequence and Topology Revealed by Molecular Dynamics Simulations"

### Data clustering

We used agglomerative hierarchical clustering with complete linkage to identify the principal configurations of the $3_1$-knotted SP along the unfolding by translocation pathways obtained using axial pulling forces of different average magnitude. Specifically, we compared trajectories at the average force of 300 pN, the default one for $3_1$-knotted SP and a lower force of 200 pN. The latter is typically too weak to induce complete translocations over the simulated timescales and thus we compare it to the 300 pN trajectories that did not result in complete translocation. For our analysis, we applied the Euclidean metric to datasets of the fraction of native contacts $Q_N$ and translocated fraction $x$. The appropriate number of clusters, 7 for 200 pN trajectories and 8 for 300 pN trajectories, was determined using the average silhouette,[1] Caliński–Harabasz[2] and Davies–Bouldin[3] scores.

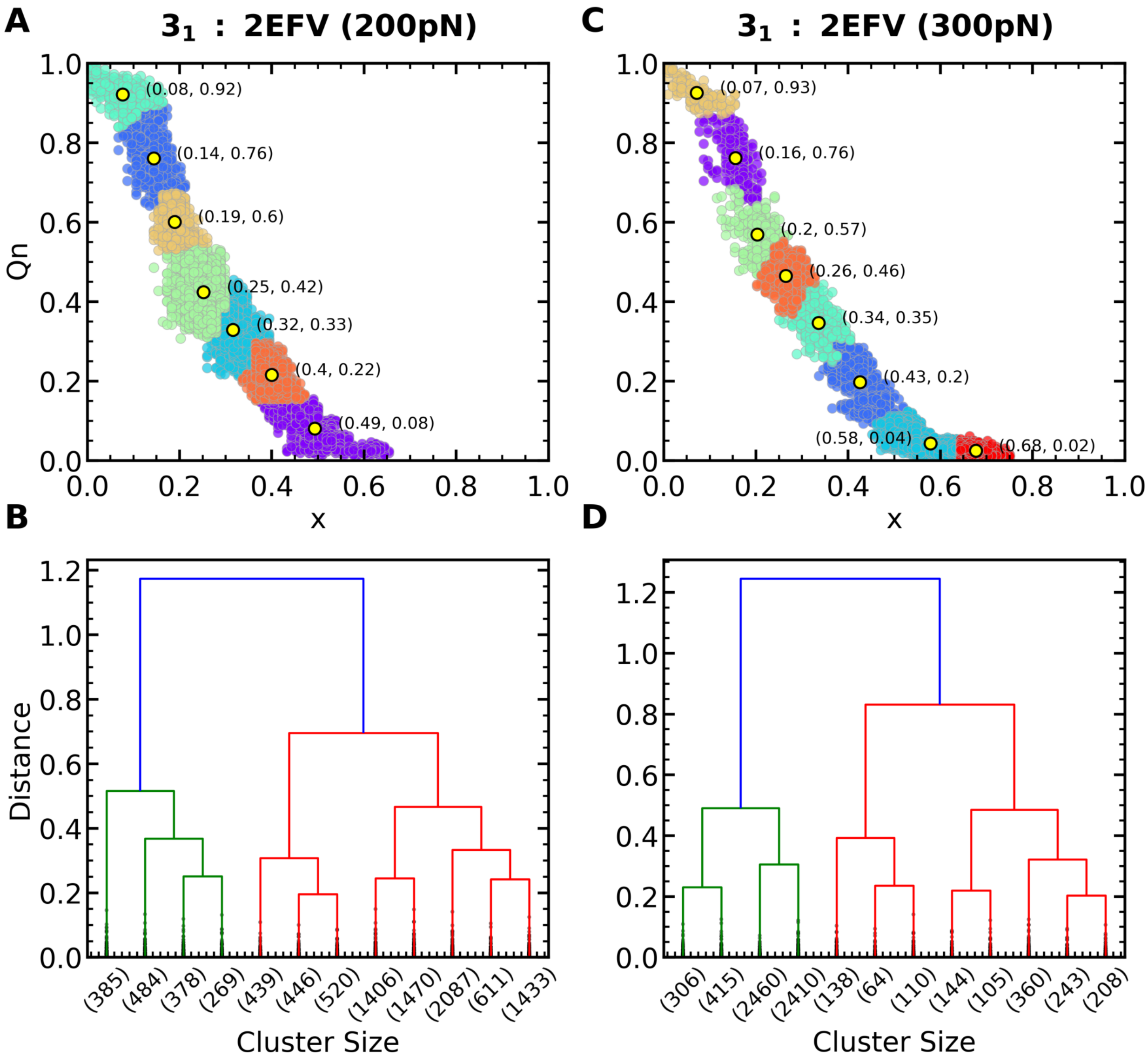


Figure S1: Clustering analysis of unfolding and translocation pathways using different pulling forces. Principal clusters and truncated dendrograms (including the top 12 leaves) of datasets of $Q_N$ and translocated fraction $x$ in pathways obtained using axial pulling forces with average magnitudes of (A)–(B) 200 pN and (C)–(D) 300 pN. Centroids of clusters are indicated using yellow dots.

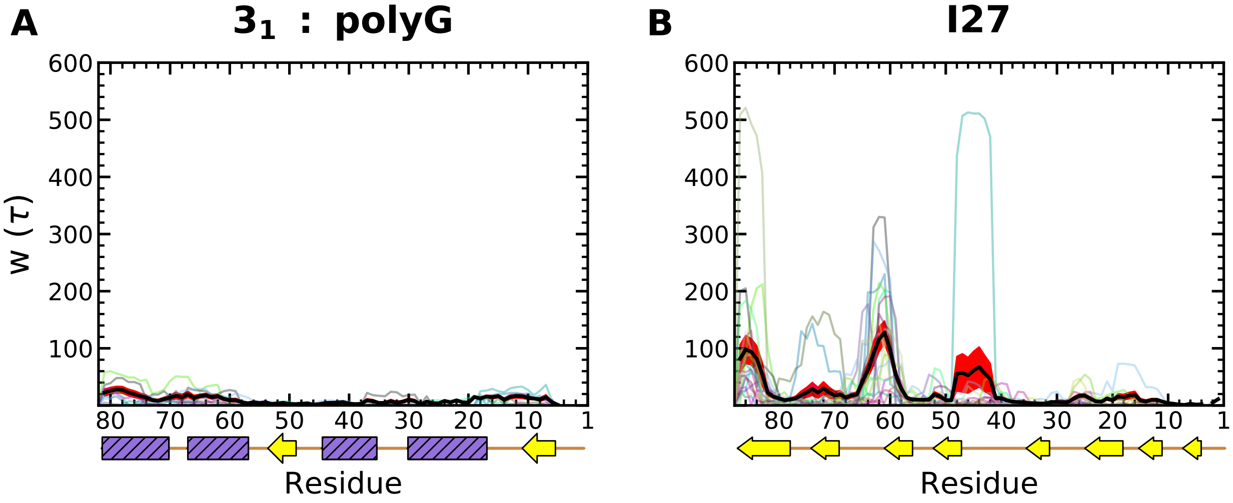


Figure S2: Waiting times per residue in translocation pathways of (A) the polyG $3_1$–knotted SP and (B) I27. Thin curves indicate individual trajectories and the thick black curves show the averaged values. The red bands highlight the standard errors.

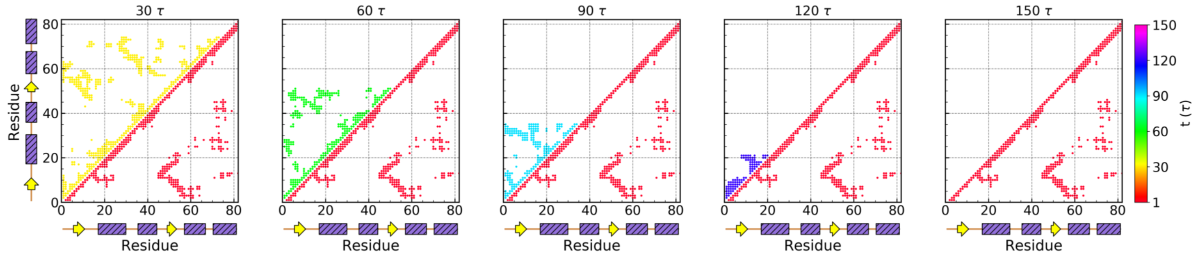


Figure S3: Contact map of the $3_1$–knotted polyG SP. The time evolution of the contact map of the SP is shown in the upper triangles using time–dependent colors. The native contact map (red) is indicated in lower triangles.

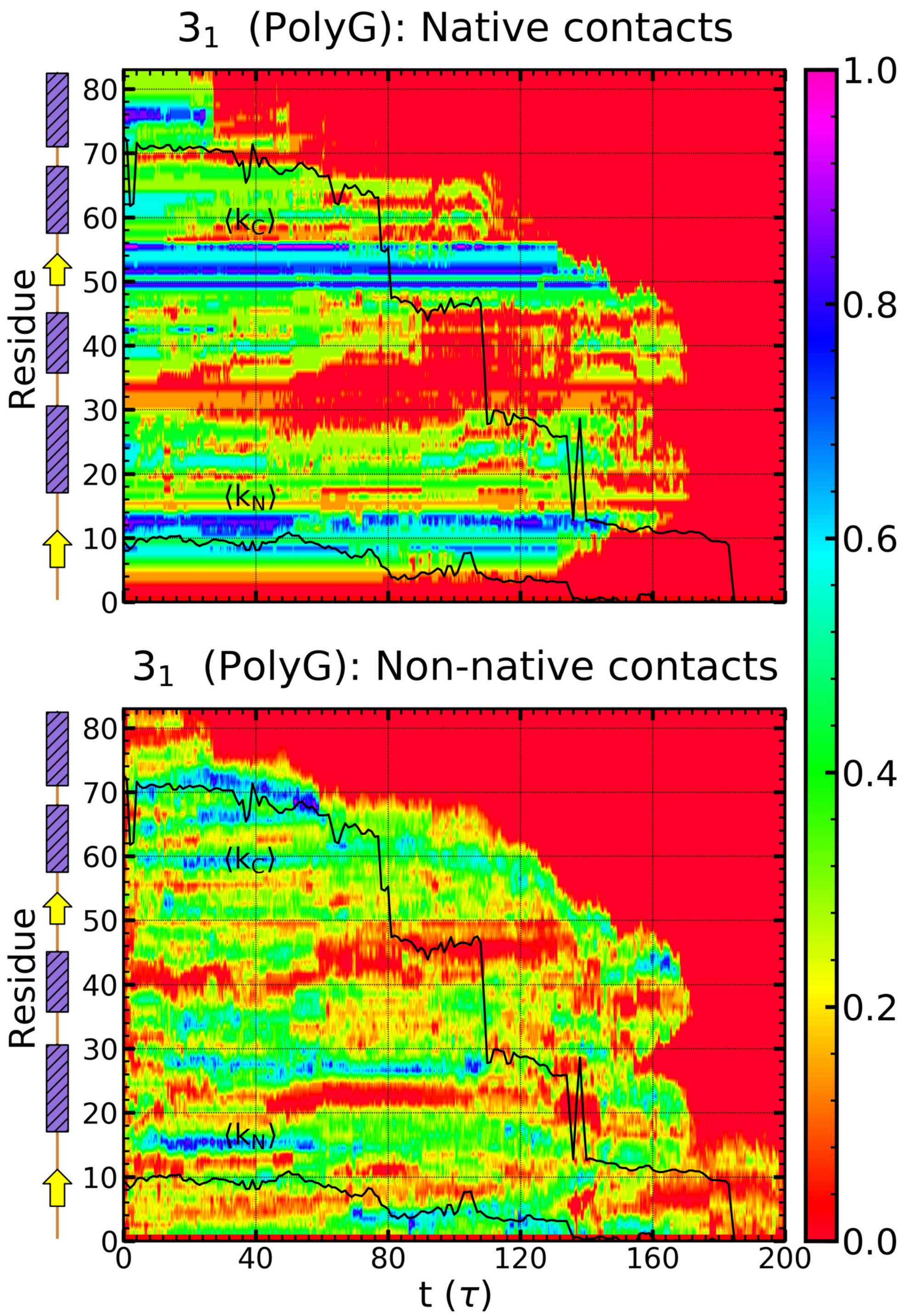


Figure S4: Time evolution of the fraction of native contacts $Q_N$ and non–native contacts $f_{NN}$ formed by secondary structural elements of the $3_1$–knotted polyG SP. The time–dependent average knot boundaries are indicated with black curves.

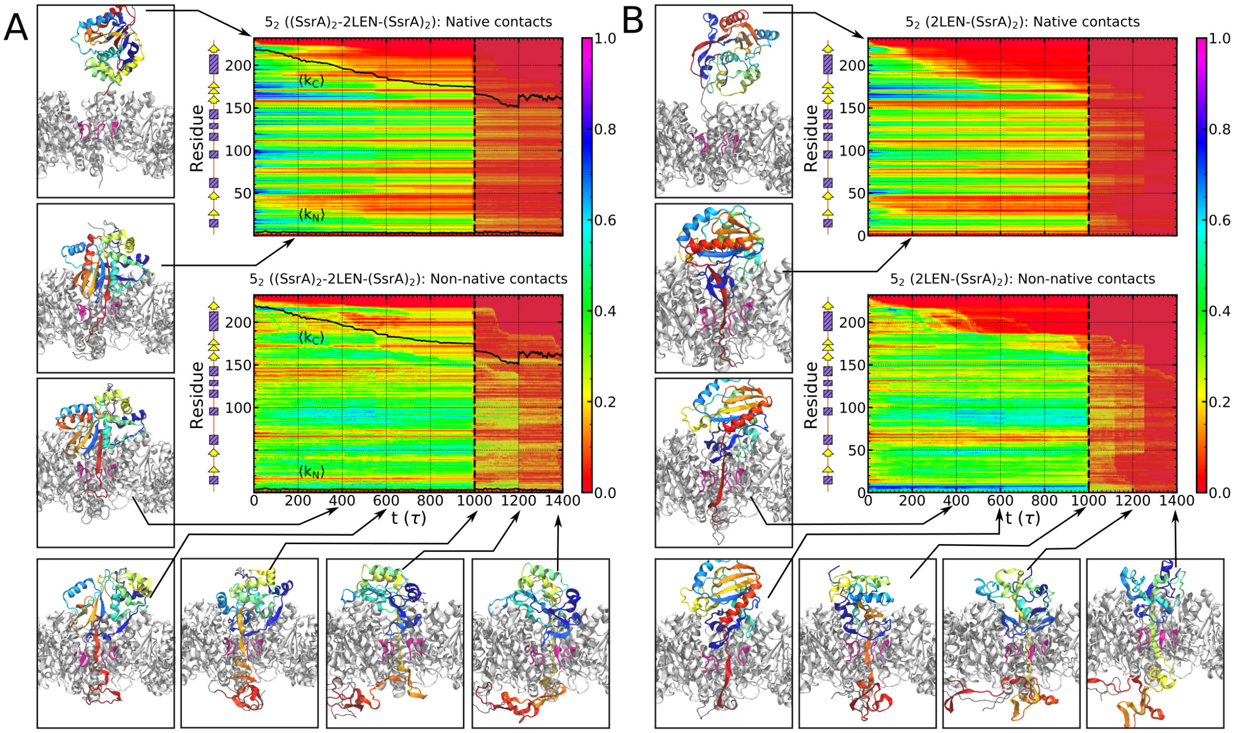


Figure S5: Time evolution of the fraction of native contacts ($Q_N$, upper panels) and of the fraction of non–native contacts ($f_{NN}$, lower panels) of individual residues in unfolding and translocation of (A) the $(SsrA)_2$–2LEN–$(SsrA)_2$ and (B) the 2LEN–$(SsrA)_2$ SPs in the C–N direction mediated by ClpYΔI. Also shown are representative configurations of the ClpYΔI–SP system during simulation trajectories and the average knot boundaries (black curves). Shaded regions correspond to fast–forwarding of selected trajectories for $1000\tau < t < 1400\tau$ using axial pulling forces with average magnitude of 600 pN.

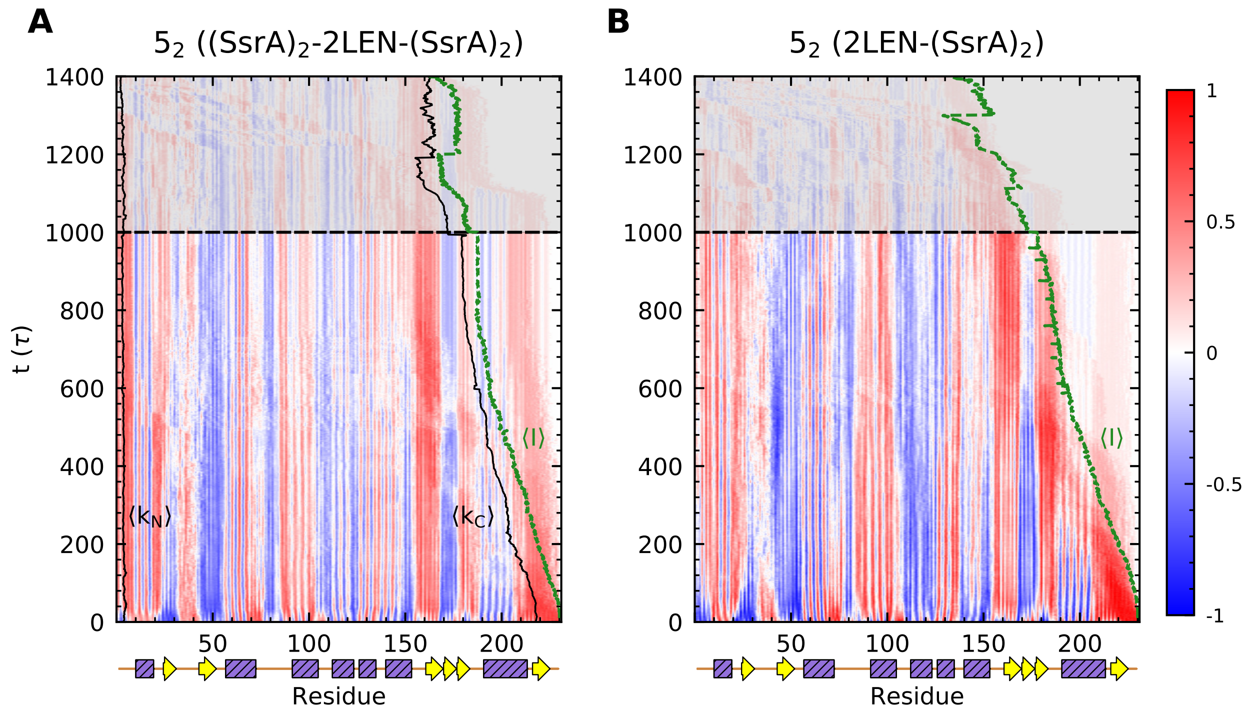


Figure S6: Tension propagation within the $5_2$–knotted SPs processed in the C–N direction. Average time–dependent orientation of virtual $C_\alpha$–$C_\alpha$ bonds involving successive amino acids, $\cos\theta$, is indicated with respect to the axial force direction for (A) the $(SsrA)_2$–2LEN–$(SsrA)_2$ SP, for which the knot persists and (B) the 2LEN–$(SsrA)_2$ SP, for which the knot is removed. The average knot boundaries are also indicated (black curves). Shaded regions correspond to fast–forwarding of selected trajectories for $1000\tau < t < 1400\tau$ using axial pulling forces with average magnitude of 600 pN.
